# Active foam dynamics of tissue spheroid fusion

**DOI:** 10.1101/2024.08.30.610303

**Authors:** Steven Ongenae, Hanna Svitina, Tom E. R. Belpaire, Jef Vangheel, Tobie Martens, Pieter Vanden Berghe, Ioannis Papantoniou, Bart Smeets

## Abstract

Three-dimensional tissue spheroids are a key building block in biofabrication, yet the link between their material properties and active mechanics of individual cells is not fully understood. We study the material properties of small spheroids of human periosteum-derived cells as they effect spheroid fusion, an elementary operation for constructing large tissue structures. We use two-photon confocal microscopy to measure cell-cell tension and individual cell motility throughout fusion. Cytoskeletal inhibition through Y-27632 (ROCKi) results in more granular tissues with decreased cell rearrangements, but accelerated fusion. Further reducing cell contractility with blebbistatin and ROCKi increases tissue granularity, decreases rearrangements, and slows down fusion. In all conditions, complete fusion is associated with frequent cellular rearrangements. Using a novel computational model that represents tissue material as an active cellular foam, with cells depicted as viscous shells with interfacial tension and persistent, random motility, we construct a phase diagram of spheroid fusion in function of relative cell-cell tension and cell motility. Our results reveal a close relationship between microscopic tissue fluidity and the visco-elastic properties of spheroid fusion. Additionally, we find that cell-cell friction promotes arrested fusion by inducing jamming through a distinct physical mechanism. Combined, our findings offer a framework for understanding spheroid fusion dynamics that can aid in the robust generation of large tissue constructs for regenerative medicine.

## Introduction

Tissue spheroids are small self-organized cell aggregates resembling tissues or micro-tumors. They offer a versatile platform for exploring cellular behavior in a 3D context [1] and play an important role in drug discovery, disease modeling, and various applications in regenerative medicine [2]. Their small size effectively overcomes diffusion limitations encountered in larger tissues, while their self-organized 3D micro-environment serves as an optimal setting for recapitulating various developmental pathways, including chondrogenic differentiation [3, 4, 5]. In tissue engineering, they serve as building blocks for tissue grafts that are suitable for transplantation [6, 7].

An essential step in the assembly of tissue spheroids into larger tissues is the fusion of two spheroids. Fusion not only serves as the unit operation for the fabrication of larger tissue constructs, it also offers a controlled, *in vitro* setting to study the properties of cellular condensations in development [8]. Spheroid fusion can be modeled as the coalescence of two viscous droplets driven by surface tension, forming a single spherical aggregate by minimizing free surface energy [9]. In this model, fusion dynamics depend on spheroid size, material viscosity, and effective surface tension [10]. Spheroid fusion may also arrest, preserving aspects of the original spheroid shape and spatial structure in large tissue constructs [11, 12]. Arrested dynamics can be modeled by the buildup of elastic energy during fusion [13].

At the cell scale, this apparent elastic energy stems from the inability of cells to undergo topological rearrangements, where cells switch neighbors [14]. As such, it relates to the jamming transition, in which cells effectively become caged in their local neighborhood due to energy barriers resulting from cell-cell adhesion, or from the rigidity of the nucleus, the cell or the extracellular matrix [15]. External forces, but also the presence of internal active forces due to motility, can help overcome these energy barriers to recover tissue fluidity [16].

Interest in rigidity transitions has grown due to their crucial role in tissue transformations where phenotypic transitions are enabled, such as those encountered in embryonic development and disease [17, 18, 19, 20]. These transitions allow tissues to rapidly shift between viscous and elastic behaviors during development [21, 22]. Still, the implications of these transitions for tissue engineering and biofabrication applications remain largely unexplored. While it is anticipated that cell motility influences both fusion degree and timescale, there lacks an overarching theory linking cellular-scale fluidized or jammed states to tissue-scale visco-elastic and rheological properties [23]. Computational models such as the vertex [24] and self-propelled Voronoi model [16] have greatly advanced our understanding of the jamming transition in biology. Recent expansions of these models also include inter-cellular pores in a dynamic ‘active foam’ [25]. Moreover, the representation of tissues as cellular foam has emerged as a powerful paradigm for force inference based on shape in 3D cell aggregates [26, 27]. Still, current foam models lack the capability to capture the active, 3D dynamics of tissue spheroids.

This study investigates the fusion dynamics of small spheroids of human periosteum-derived cells (hPDCs) used in bottom-up bone and cartilage engineering [3]. hPDCs are instrumental for the formation of the fracture callus, a transient cartilaginous tissue that enables fracture healing [28]. For skeletal tissue engineering, *in vitro* multicellular condensations of these cells have been engineered to replicate skeletal developmental cascades [28, 3]. By combining microscopy experiments, visco-elastic theory and simulations of a novel active foam model, we explore the link between fusion dynamics and the microscopic fluidized or jammed state of the tissue material. Additionally, we uncover how tissue fluidity scales with active mechanical properties at the cellular scale.

## Results

### Small tissue spheroids display visco-elastic fusion dynamics

To obtain statistically reliable data on fusion dynamics, we conducted high-throughput time-lapse microscopy of fusing spheroids containing approximately 100 hPDCs in chondrogenic combination medium (C8), comprising Y-27632 Rho kinase inhibitor (ROCKi) and TGF-*β* 1, along with other components (Fig. 1a). ROCKi, an inhibitor of the Rho-ROCK signaling pathway, is known for suppressing actin polymerization [29] and has been previously shown to qualitatively influence aggregation dynamics [30] and enhance fusion capacity [3]. TGF-*β*, an epithelial-to-mesenchymal transition inducer, is recognized for its role in promoting chondrogenesis in mesenchymal cells [31, 32]. We characterized the fusion angle *θ* based on the analogy between spheroid fusion and the coalescence of two spherical droplets (Fig. 1b). Elimination of either ROCKi or TGF-*β* 1 resulted in deceleration of fusion (Fig. 1c). In an effort to diminish cellular force generation, we enriched the medium with a higher concentration (100 µM) of ROCKi and introduced 20 µM of blebbistatin, an inhibitor of non-muscle myosin II that reduces cell contractility. These conditions led to a distinct slowdown in fusion dynamics. Yet, viability was maintained and fusion persisted. To quantify these differences, we fitted individual fusion curves with a solution to the governing equation for fusion of visco-elastic droplets [13],

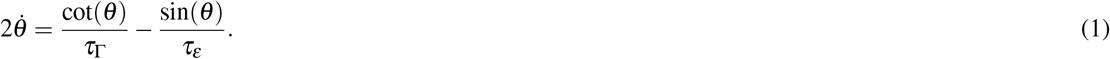

**Fig. 1:**
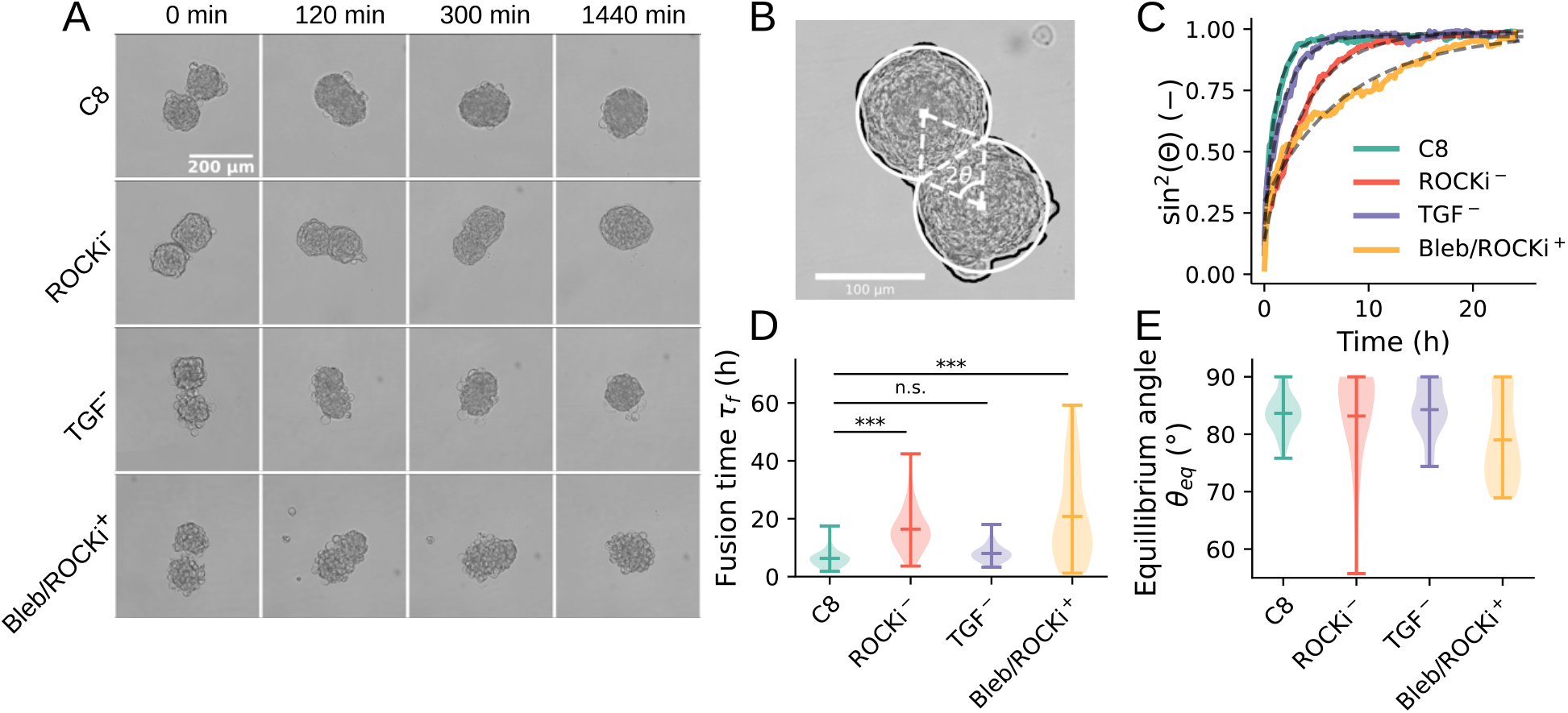
Fusion dynamics of hPDC spheroids in 4 conditions: C8 medium (C8), C8 w/o Y-27632 Rho kinase inhibitor (ROCKi^−^), C8 w/o TGF-*β* 1 (TGF^−^) and C8 + 100 µm Y-27632 Rho kinase inhibitor and 25 µm blebbistatin (Bleb/ROCKi^+^). **A**: Bright-field microscopy snapshots at 0, 120, 300 and 1440 min after start of fusion. **B**: Illustration of the fusion angle *θ*, with fitted circular contours for each spheroid. **C**: Temporal evolution of the average fusion angle for 4 treatment conditions, together with the fit of Eq. (1). Data acquired from a single experiment, *n* = 7, 10, 8, 9, for resp. C8, ROCKi^−^, TGF^−^, Bleb/ROCKi^+^ **D**: Fusion timescale *τ*_*f*_ and **E**: Equilibrium fusion angle *θ*_*eq*_ for 4 treatment conditions. Data is pooled from different experiments, see Supplementary Fig. 1, with *n* = 38, 46, 44, 22 for resp. C8, ROCKi^−^, TGF^−^, Bleb/ROCKi^+^. Significance with Dunnett’s test comparing treatments versus C8 (n.s: *p >* 0.05, ‘***’: *p <* 0.001)

Here, *τ*_Γ_ = *R*_0_*η*_*t*_*/*Γ is the visco-capillary time and *τ*_*ε*_ = 4*η*_*t*_*/G*^*’*^ is the visco-elastic with initial spheroid radius *R*_0_, tissue viscosity *η*_*t*_, tissue surface tension Γ, and shear modulus *G*^*’*^. This model fits the data well in all studied conditions (Fig. 1c and Supplementary Fig. 1-2). We extracted two key fusion parameters: the equilibrium fusion angle *θ*_*eq*_, with sec(*θ*_*eq*_) − cos(*θ*_*eq*_) = 4Γ*/*(*G*^*’*^*R*_0_), and the fusion time-scale 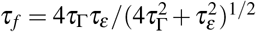..Spheroids cultured in absence of ROCKi exhibited significantly longer fusion times compared to the baseline C8 condition (Fig. 1d). Moreover, the fusion time was significantly increased for spheroids with blebbistatin and elevated levels of ROCKi. Despite large variations, most conditions exhibit nearly complete coalescence, although equilibrium fusion angles less than 90^°^ are observed for treatment with both blebbistatin and elevated levels of ROCKi (Fig. 1e).

### Micro-structure and dynamics reveal distinct cell mechanics and motility

Subsequently, we investigated the relation between fusion dynamics and micro-structure and dynamics at the cell scale. Cell-cell interfacial tension governs tissue surface tension (Γ) at the spheroid scale [33, 34], which in turn sets the fusion time-scale *τ*_*f*_. In a foam, the relative interfacial tension is discernible in the contact angle at triple junctions [26]. We used confocal microscopy imaging to reveal the shape and internal structure of individual spheroids in the aforementioned conditions. In absence of ROCK inhibition, both nuclei and actin structures exhibited increased orientation in the circumferential direction compared to other conditions, which displayed a more isotropic distribution (Fig. 2a). Additionally, significant variations in spheroid shape granularity were observed between conditions. To quantify these differences in granularity, we analyzed the angular distribution of curvature in cells at the periphery of the spheroid (Fig. 2b). From this analysis, we define the granular shape index as GSI = *r*(*κ*−1*/R*), where *r* is the mean cell radius, *R* the spheroid radius, and *κ* the median sector curvature (Fig. 2c). The GSI is zero for a perfectly smooth spherical aggregate and approaches 1 for a loosely packed, granular cell arrangement with radius *r*. Through simulation of a static foam with surface tension *γ* and adhesive tension *w*, we found that, for large enough aggregates (*>* 50 cells), the GSI well approximates the relative cell-cell tension GSI ≈ *α* = 1 − *w/γ* (Extended Data Fig. 1). Unlike methods that estimate *α* using contact angles [35, 21], this approach avoids the potentially error-prone task of precisely delineating cell-cell interfaces, especially in smooth spherical aggregates. The GSI was approximately 0.5 for C8 both with and without TGF-*β* 1, dropped significantly to around 0.25 in the absence of ROCKi, and increased significantly, approaching 0.75, with blebbistatin and elevated ROCKi levels (Fig. 2d). Thus, while cell-cell tension governs tissue surface tension, the observation of faster fusion times in C8 suggests that there is no singular connection between cell-cell tension and fusion time.

**Fig. 2:**
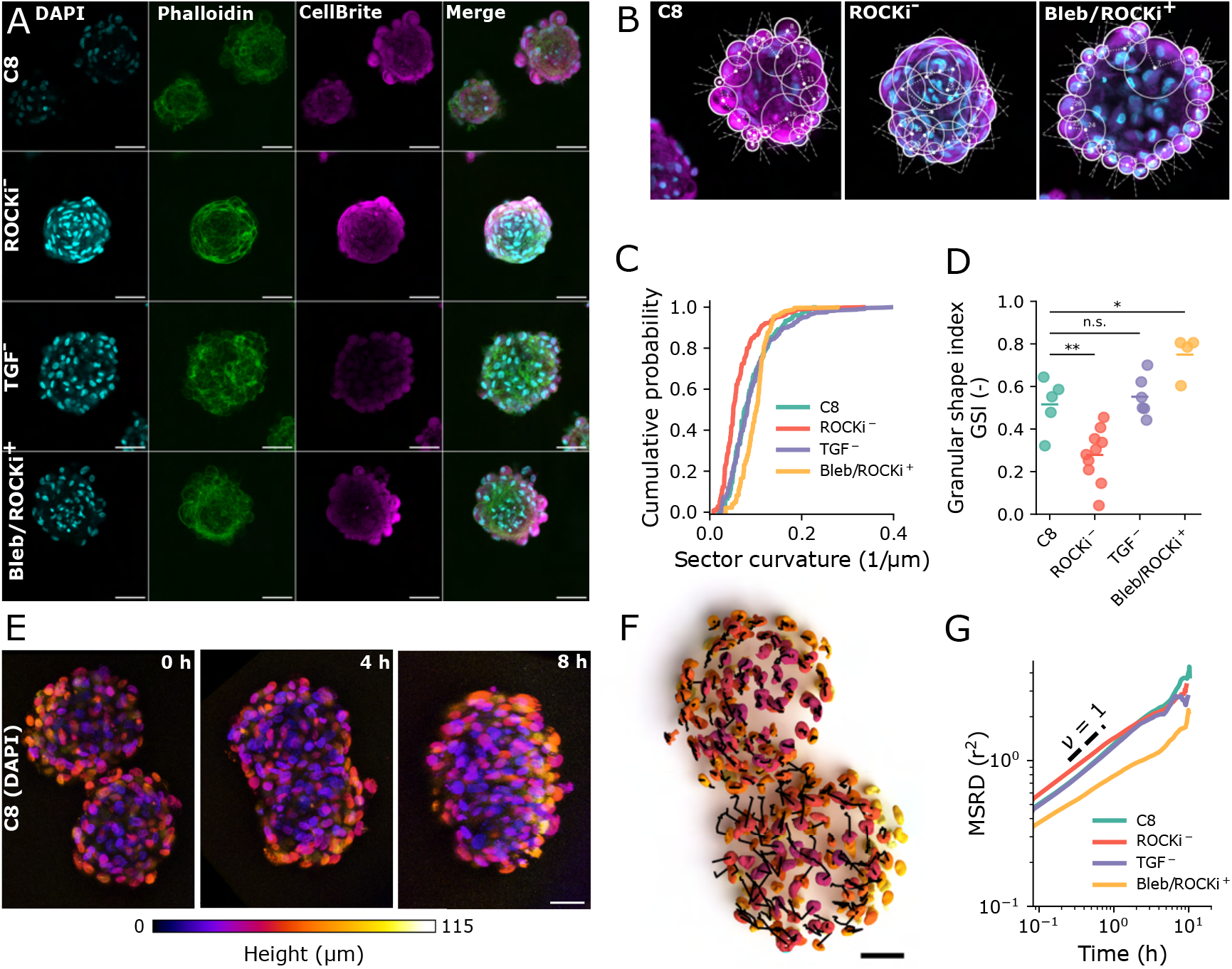
Spheroid shape and cell motility. **A**: Confocal microscopy slices of spheroids in conditions C8, ROCKi^−^, TGF^−^ and Bleb/ROCKi^+^, with stainings DAPI (nucleus), Phalloidin (actin) and CellBrite (membrane). **B**: Circular approximation of spheroid periphery, showing contour radii and contact angles for 4 conditions. **C**: Cumulative distribution of contour curvature, extracted from contour circles, in 4 conditions. Data is pooled with *n* = 5, 10, 6 and 4 for resp. ROCKi^−^, TGF^−^ and Bleb/ROCKi^+^. **D**: Granular Shape Index (GSI) for 4 conditions. Data is from individual spheroids as used in C. Significance with Dunnett’s test, comparing treatments versus C8 (n.s: *p >* 0.05, *: *p* = 0.016, **: *p* = 0.003). **E**: Time-lapse of cell nuclei (DAPI staining) from two-photon microscopy imaging colored by height. **F**: Cell trajectories (black) traced from time-lapse confocal microscopy imaging of tissue spheroid fusion (C8 condition). **G**: Time-averaged mean squared relative displacement (MSRD) for fusing spheroids in 4 conditions. The dashed line indicates diffusive behavior (exponent *ν* = 1). All scale-bars indicate 50 µm.

We then explored the impact of active cell motility on fusion dynamics. Using time-lapse two-photon microscopy, we observed fusing spheroids across the 4 studied conditions (Fig. 2e). Cell trajectories were determined by tracking nucleus positions, and we computed the time-averaged mean squared relative displacement (MSRD) for all cell pairs within the Delaunay graph of their positions (Fig. 2f). These relative displacements signify cellular rearrangements that are expected to occur frequently in a fluidized state, while being nearly absent in a solid, jammed tissue state [25]. Relative displacements were greater in absence of ROCKi (Fig. 2g), consistent with the known function of Rho kinase inhibitor in suppressing cytoskeletal rearrangements and subsequently reducing cell motility [36]. Additionally, relative displacements markedly decreased with blebbistatin and increased ROCKi, indicating reduced cellular rearrangements. In all other conditions, the lag time for topological rearrangements, signified by MSRD *> r*^2^, was under one hour, confirming that the tissue material exhibited fluid-like behavior within the experimental timescale. Interestingly, despite decreased relative cell-cell tension and cell motility, fusion proceeded notably slower in absence of ROCKi. Upon elimination of TGF-*β* 1, granularity and relative displacements showed no significant differences from C8, consistent with similar fusion dynamics.

### Active foam model captures observed fusion dynamics

To simulate the mechanical and rheological properties within multicellular aggregates, we implemented a 3D active foam model where cells are represented as discrete, deformable entities with foam-like mechanics. Cells are modeled as pressurized cortical shells with average radius *r*, planar viscosity *η* and free surface tension *γ*. They interact via adhesive tension *w*, representing the reduction of interfacial tension at the cell-cell interface [37], and migrate via persistent, random protrusion forces (Fig. 3a). The key parameters governing fusion are: i) the relative cell-cell tension *α* =−1 *w/γ*, ii) the relative active pressure *p*_*a*_*r/*2*γ*, iii) the persistence time *τ*_*p*_, and iv) the hydrodynamic length *L*_*η*_ = (*η/ξ*)^1*/*2^ with cell-cell friction constant *ξ* [38]. Observing diffusive behavior in the mean-squared relative displacements for all lag times (Fig. 2g), we infer that *τ*_*p*_≪*τ*_*f*_, and thus set *τ*_*p*_ = 10 min. Additionally, we first assume that the viscous length scale is significant, implying low cell-cell friction, setting *L*_*e*_*ta/r* = 0.5. Through modulation of cell-cell tension *α* and subsequent condensation and equilibration, we produced virtual tissue spheroids containing 100 cells with varying levels of granularity, as shown in Fig. 3b. When brought into contact, the simulated spheroids underwent spontaneous fusion (Fig. 3c). In line with our experimental observations, fusion dynamics closely align with the analytical solution derived from the governing equation of visco-elastic fusion, Eq. (1), both when varying cell-cell tension (Fig. 3d) and cell motility (Fig. 3e). This permits the estimation of the equilibrium fusion angle and the fusion time scale from simulations. When varying both 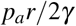 and *α*, we found that complete fusion (*θ*_*eq*_ → 90^°^) requires high motility and low relative cell-cell tension, i.e., high adhesive tension (Fig. 3f). Conversely, arrested coalescence occurred in conditions of low motility and/or high cell-cell tension. Fusion time-scale first increases with *p*_*a*_ at low motility, while it decreases with *p*_*a*_ at higher levels of motility (Fig. 3g).

**Fig. 3:**
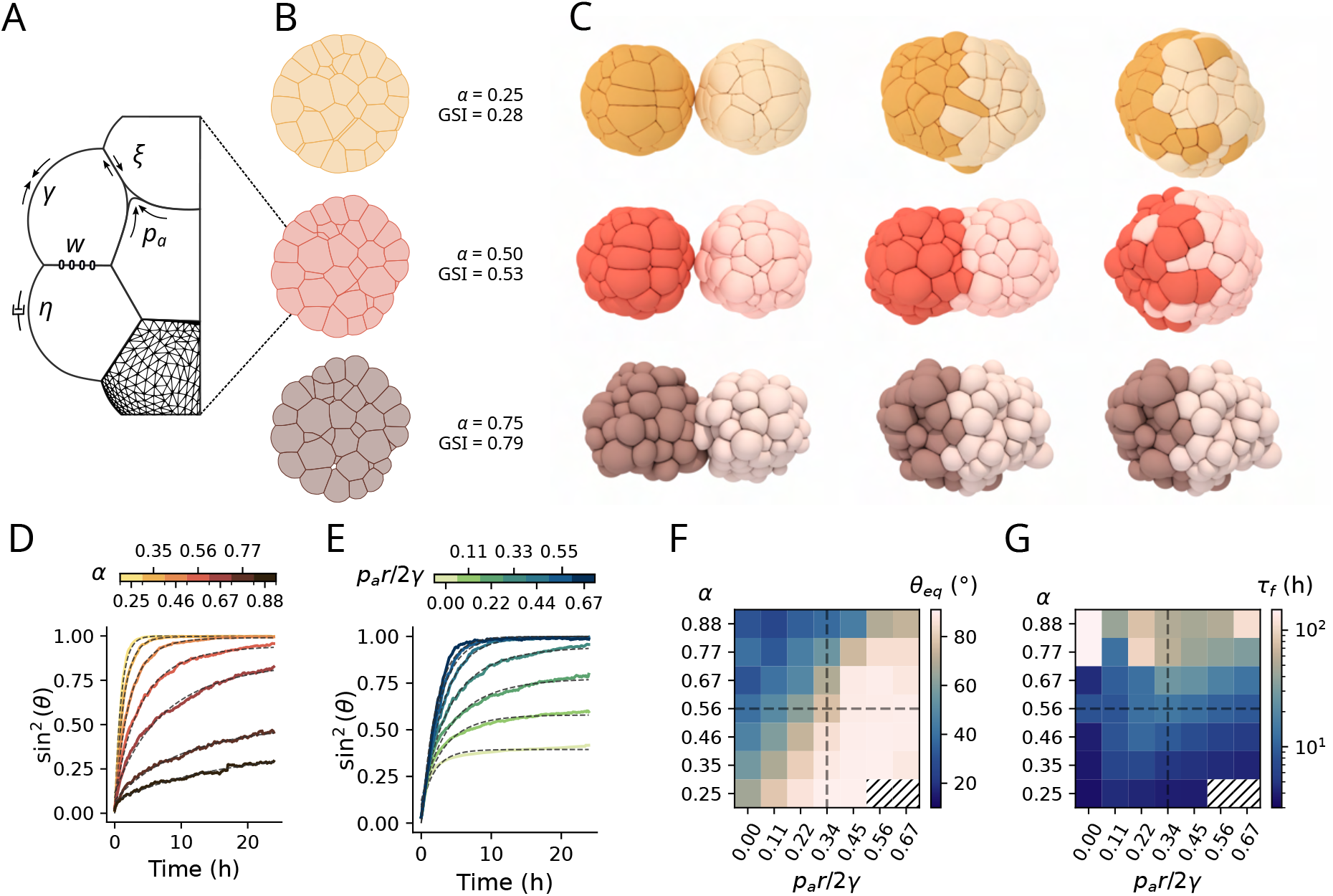
Active foam simulation of spheroid fusion. **A**: Model overview, showing cells with surface tension *γ* at the cell-medium interface, viscosity *η* and active pressure *p*_*a*_, interacting with adhesive tension *w* and cell-cell friction *ξ*. **B**: Cross-sections of simulated spheroids at cell-cell tension *α* = [0.25, 0.50, 0.75], together with the GSI obtained as in the experiments (Fig. 2d). **C**: Temporal snapshots of simulated fusion for *α* = 0.25, 0.5 and 0.75 at 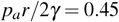. **D**: Temporal evolution of the average simulated fusion angle at fixed active pressure 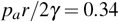 and varying *α*, together with fitted dynamics using Eq. (1). **E**: Temporal evolution of the average simulated fusion angle at fixed *α* = 0.56 and varying *p*_*a*_, together with fitted dynamics. **F**: Equilibrium fusion angle *θ*_*eq*_ for varying *α* and *p*_*a*_. **G**: Fusion time-scale *τ*_*f*_ for varying *α* and *p*_*a*_. Vertical and horizontal dashed lines in (f) and (g) indicate the parameter range for resp. panels (d) and (e). Hatched data points in (f,g) show simulations that failed to converge at high motility and low *α* due to highly negative surface tension at the cell front (*p*_*a*_*r/*2*γ ≫ α*).

### Fusion outcome reflects microscopic tissue fluidity

By tracking simulated cells, we then measured the time-averaged mean-squared relative displacement, mirroring the experimental procedure (Fig. 4a-b). As cellular motility increases and cell-cell tension decreases, we observe a transition in the mean-squared relative displacement. At low motility, it shows a sub-diffusive plateau at intermediate lag times, indicating a jammed configuration [40], which vanishes at higher cell motility, suggesting a fluidized state. We then analyzed cell rearrangements by counting unique contacts during fusion, a measure indicating fluidity (Fig. 3c). We found that the critical active pressure to fluidize the tissue material scales as *p*_*a*,crit_ ∝ *α*^−1*/*2^. In the studied range of cell-cell tension *α*, the tissue is mostly confluent [21]. In this regime, the ability of cells to deform is expected to drive topological rearrangements [24]. The role of cell deformation in tissue fluidity is evident in the 3D shape index *S/V* ^2*/*3^, with cell area *S* and volume *V* (Fig. 3k). The critical shape index of 5.41, which was previously derived from a 3D Voronoi model [39], effectively marks the jamming transition at high levels of cell motility. Through the variation of parameters *α, p*_*a*_, *τ*_*p*_, and *γ*, we identified a dimensionless activity parameter 𝒜 that collapses the scaling of cell rearrangements,

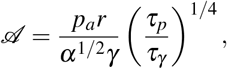

with *τ*_*γ*_ = *η/γ* (Fig. 4e and Extended Data Fig. 2-3). For small spheroids, the effect of spheroid size on tissue fluidity is weak (Extended Data Fig. 4 and Supplementary Fig. 3). Interestingly, this scaling deviates from the behavior of active Brownian particle models or self-propelled Voronoi models with self-propulsion velocity *v*_0_, where an effective temperature 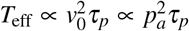 governs the glass transition [16]. This suggests that i) *τ*_*p*_ is large enough so that rearrangements cannot be assumed to be only small and localized, but can be collective [16] and/or ii) finite size effects and the influence of cell motility on tissue shape may qualitatively alter rearrangement dynamics. In our simulations, the degree of coalescence is closely tied to microscopic tissue fluidity, as i s evident from the comparison between equilibrium fusion angle and the mean squared displacement for a 60 min lag time — an experimentally accessible parameter that follows the normalized unique contact count (Fig. 3f). Jamming coincides with arrested coalescence, where *θ*_*eq*_ and, correspondingly, the tissue viscoelastic timescale *τ*_*ε*_, sharply increase with activity. As the cellular material fluidizes, coalescence becomes complete, and *τ* _*ε*_→∞. At high activity levels, the equilibrium fusion angle decreases again due to active shape fluctuations in the aggregate. When mapped onto t his c hart, our experimental observations follow this general pattern. Interestingly, they are located close to the fluid-to-solid t ransition i n the corner, which is traversed with blebbistatin and elevated levels of ROCKi. Rearrangements due to cell activity also enhance mixing during fusion. At low activity, cells from separate spheroids maintain distinct interfaces. However, increased activity leads to intermixing of cells from different spheroids in the timescale of fusion (Fig. 4g), resulting in the loss of distinct boundaries between the original tissues (Fig. 3c).

**Fig. 4:**
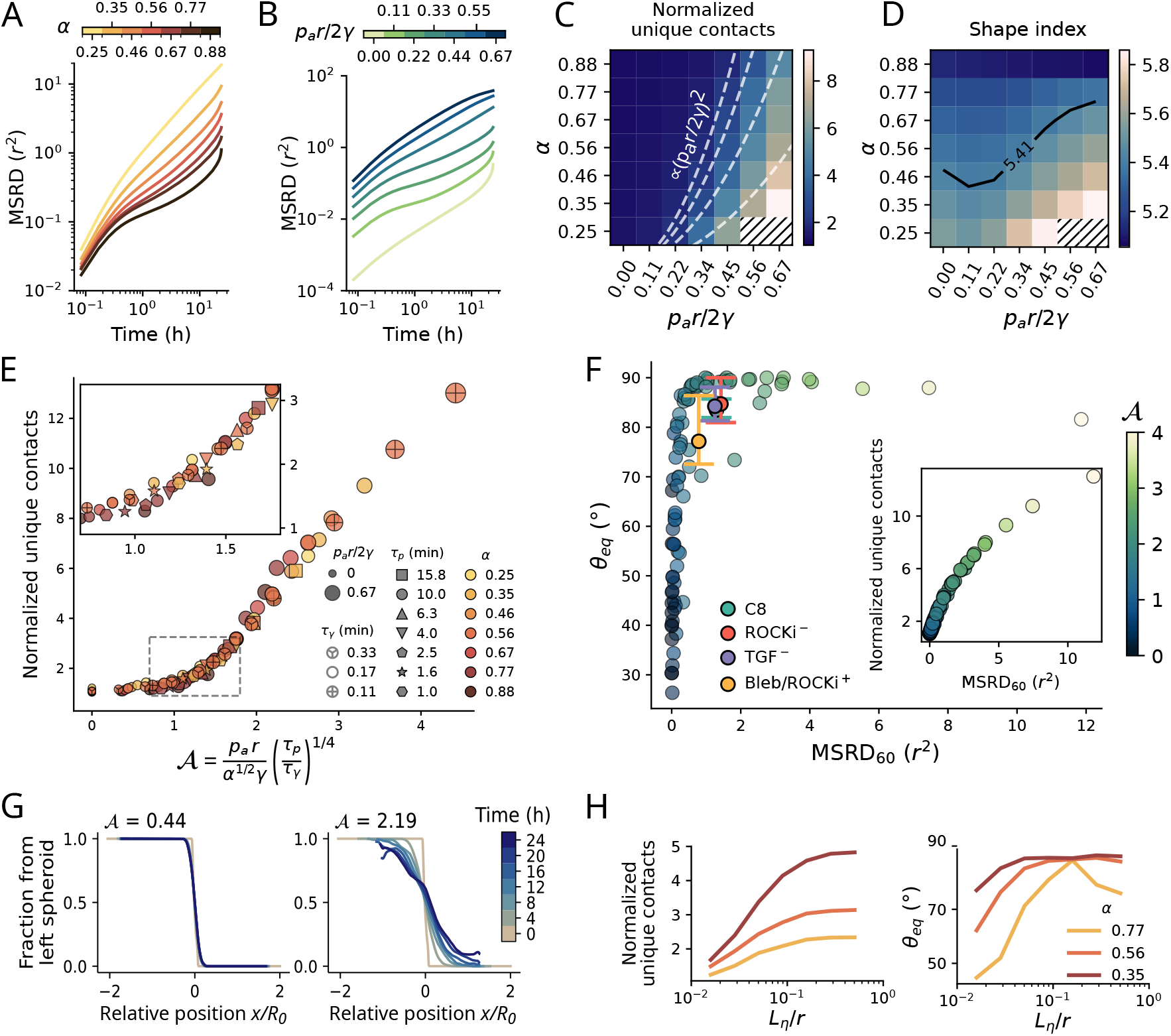
Tissue fluidity during spheroid fusion. **A**: Mean-squared relative displacements (MSRD) for varying cell-cell tension *α* at fixed motility 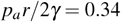.**B**: MSRD for varying 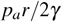 at fixed *α* = 0.56. **C**: Normalized unique contact number (total number of unique contacts during fusion divided by average number of contacts) for varying *α* and *p*_*a*_. Dashed lines show the scaling 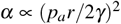.**D**: 3D shape index *S/V* ^2*/*3^ with cell surface area *S* and volume *V* for varying *α* and *p*_*a*_. The black line shows the iso-contour of shape index 5.41 [39]. Hatched data points in (c,d) show simulations that failed to converge at high motility and low *α*. **E**: Normalized unique contacts for varying *p*_*a*_ (marker size), *τ*_*γ*_ (marker hatching), *τ*_*p*_ (marker shape) and *α* (marker color), see further Supplementary Fig. 3-4, showing collapse of data points for 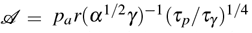. **F**: Fitted equilibrium fusion angle *θ*_*eq*_ versus mean squared relative displacement for a lag time of 60 min (MSRD_60_) for combined simulation data, where color scale shows activity 𝒜, and compared to experimental observations in 4 conditions. Inset: MSRD_60_ follows normalized unique contacts in simulations. **G**: Temporal evolution of aggregate mixing for low (left) and high (right) levels of cell activity. **H**: Normalized unique contact number (left) and equilibrium fusion angle (right) in function of *L*_*η*_ for varying *α*.

The active foam model treats cells as discrete entities capable of relative sliding, allowing us to evaluate the impact of cell-cell friction on spheroid fusion and fluidity. At slow timescales, the dynamic binding and unbinding of adhesive ligands leads to an effective wet friction, which is parameterized in our model by the hydrodynamic length-scale *L*_*η*_ [41]. Hence, increasing the bond lifetime of adhesive ligands corresponds to a decrease of *L*_*η*_. As *L*_*η*_ decreases, the normalized unique contact count and the equilibrium fusion angle notably reduce (Fig. 4h), indicating that cell-cell friction facilitates arrested coalescence. Interestingly, jamming also occurs for elongated cells with a high shape index, suggesting that cell-cell friction offers a distinct mechanism to induce jamming compared to shape-dependent rigidity transitions [24, 39] (Extended Data Fig. 5).

## Discussion

Small tissue spheroids are well-suited for examining tissue rheology in 3D due to their well-defined boundaries, compact size allowing for precise measurement of individual cell shape and dynamics, and the ability to study dynamic processes such as fusion, spreading, dispersion, and aspiration. In this work, we explored the fusion dynamics of small tissue spheroids, conceptualizing them as active cellular foams. In small spheroids of hPDCs, we found that complete fusion was associated with frequent cellular rearrangements, indicating fluidization of the tissue material. Reduction of actin polymerization and cell contractility through inhibition of ROCK resulted in more granular tissue shapes an decreased cell movements, but faster fusion. Further reducing cell contractility through blebbistatin and ROCKi led to more enhanced tissue granularity and decreased cell movements, but also slowed down fusion dynamics. These results add to prior findings in chondrogenic organoids. For instance, inhibition of ROCK during aggregate formation shifted dynamics from those of a dewetting liquid to an agitated granular pile [30]. Similarly, our results show increased spheroid granularity with inhibition of ROCK. Despite this granularity, fusion dynamics remained consistent with the coalescence of continuum, visco-elastic materials, both in experiments and simulations. However, the inferred relative cell-cell tension was below the critical value for internal pore formation (*α*≈0.875 [21]), indicating that these spheroids were internally confluent, unlike during aggregate formation.

To clarify the interplay between single cell properties and tissue dynamics, we developed a dynamic 3D active foam model with discrete, deformable cells, parameterized by relative cell-cell tension and motility. This model reproduces characteristic fusion dynamics, while capturing cell shape and relative motility. Our results show that complete fusion, corresponding to the coalescence of liquid droplets at the macroscopic level, is closely linked to tissue fluidization at the microscopic level, as illustrated in Fig. 5. We uncovered a universal scaling relationship that links tissue fluidization to relative cell-cell tension, persistence time, cortical relaxation time and active motility, through an effective activity parameter. Due to fluidization, significant variations in neighbor exchange rates are predicted between instances of near-complete coalescence. Hence, depending on cell activity, complete coalescence might involve either significant intermixing of cells from different spheroids or a quasi-stable tissue interface. This observation is relevant for biofabrication, where independent control of coalescence and intermixing may be necessary. We found that the ability of cells to elongate governs the rigidity transition, in line with vertex and Voronoi model predictions [24, 39]. Low cell-cell tension reduces energy barriers, allowing fusion akin to passive liquid behavior, while active migration forces are required to overcome these barriers at higher cell-cell tension. Classical foam and vertex models solely account for cell-cell interfaces, whereas our discrete foam model incorporates relative cell-cell sliding, which we associate with adhesive ligand or intercellular matrix turnover rates. Our results show that friction, while typically linked to transient phenomena, induces arrested fusion by promoting jamming through a distinct physical mechanism, consistent with recent phase field model predictions in cell monolayers [42]. The physical implication of cell-cell friction on tissue dynamics still needs to be explored for relevant biological mechanisms, such as cadherin turnover in epithelia [43], or the dynamic remodeling of extracellular matrix in 3D tumors and organoids [44].

**Fig. 5:**
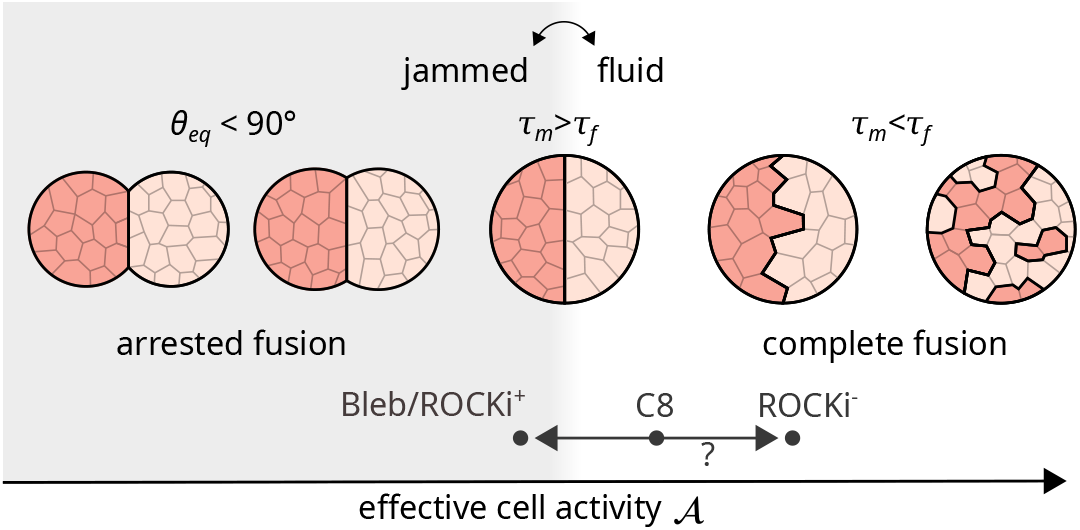
Schematic representation of active foam fusion regimes for varying the effective cell activity 𝒜. At low activity, the tissue material is jammed and arrested fusion may be observed (*θ*_*eq*_ *<* 90^°^). At high cell activity, the tissue material is fluidized and fusion is complete. Further increasing activity decreases the mixing time *τ*_*m*_ relative to the fusion time *τ*_*f*_ (Supplementary Fig. 4), resulting in homogenized cell aggregates. Based on relative cell-cell tension and apparent cell diffusion, the experimental conditions of C8, Bleb/ROCKi^+^ and ROCKi^−^ can be located on this chart. However, the increase in fusion time *τ*_*f*_ upon removal of ROCKi is not predicted by active foam simulations.

Comparison between predictions of the active foam model and our experimental observations highlights a few noteworthy similarities. Specifically, in the reference C8 chondrogenic medium, with *τ*_*f*_ ≈6.4 h, *θ*_*eq*_ ≈ 84^°^, and *α*≈0.5, our fusion simulations are compatible with experimental outcomes at intermediate levels of cell motility, Fig. 3(d-g). Furthermore, the observed increase in granularity, fusion slowdown, and decrease in relative motility after addition of both ROCKi and blebbistatin align with simulation results at elevated *α* = 0.75 and reduced or equal cell motility. This treatment, as indicated by shape index and cell rearrangements, corresponds to a transition from a fluidized state to a more jammed material, see Fig. 5. Elimination of ROCKi led to reduced granularity and increased cell movement, but also decreased fusion time. In line with this, our active foam model predicts increased relative cell movements and decreased granularity at *α*≈ 0.25 (Fig. 3b and 4a). However, while higher cell motility can slow fusion (Fig. 3g), our model cannot replicate the fusion slowdown observed with ROCKi removal. Decreasing *α* consistently speeds up fusion, and adjusting the cortical relaxation time in simulations does not match the observed increase in cell movement. This suggests the active foam model does not fully capture the biophysical changes associated with ROCKi removal, possibly due to the polarized cells with anisotropic actin structures seen in Fig. 2a. These structures, predominantly oriented in the circumferential direction, may impede radial cell movements necessary for spheroid fusion, thus slowing down fusion dynamics as they require remodeling. Future active foam models could investigate how anisotropic surface tension and cortex viscosity influences tissue structure and dynamics.

## Conclusion

We developed a framework to investigate spheroid fusion, integrating tissue-scale viscoelastic properties with cell-scale analysis of cell-cell tension and individual cell motility. We found that disrupting actin polymerization and cell contractility with Y-27632 (ROCKi) reduced cellular rearrangements, leading to more granular spheroids and slower fusion. Remarkably, eliminating ROCKi also decelerated fusion despite increased cellular rearrangements, highlighting the non-linear relationship between cell mechanics and tissue properties. Through simulations of an active foam model, we demonstrated that two distinct fusion regimes — arrested and complete — are closely linked to the tissue (un)jamming transition. Below the unjamming transition, assembloid formation is akin to colloidal sintering, with sintering degree tied to cellular activity. Beyond this transition, it resembles liquid droplet coalescence, where the mixing-to-fusion timescale ratio dictates long-term stability. These insights are crucial for the spatial and temporal assembly of large tissue constructs in biofabrication. In the future, these models can be expanded to incorporate interactions with encapsulating materials such as bioinks or the extracellular matrix. Our results on *in vitro* tissue spheroids provide insights into the physics of their *in vivo* counterparts, multicellular condensations, which play a crucial role in tissue patterning, organ formation, and cell differentiation. By taking into account the heterogeneity and asymmetries encountered *in vivo*, this theoretical framework has the potential to shed light on complex biological processes, including cell sorting, both *in vivo* and in their *in vitro* analogues.

## Methods

### Periosteal cell isolation and culture

hPDCs were isolated from periosteal biopsies of two female and four male donors aged between 11 and 17 years old. A single cell pool was created and cells were seeded at a density of 5000 cells/cm2 for expansion. Cells were incubated at 37 °C, 5% CO2 and 95% humidity until passage 9 in Dulbecco’s modified Eagle’s medium (DMEM, Life Technologies, UK) enriched with 10% fetal bovine serum (HyClone FBS, Thermo Scientific, USA) and 1% antibiotic-antimycotic (100 units/mL penicillin, 100 mg/mL streptomycin and 0.25 mg/mL amphotericin B). Expansion medium was changed every 3 days and cells were dissociated using TrypLE™ Express (Life Technologies, UK) at a confluence of 90%. For all donors informed consent was obtained and all procedures were approved (S64471) by the ethical committee for Human Medical Research (KU Leuven).

### Aggregate formation

Single-cell hPDCs suspension was seeded in U-bottom Ultra-low Attachment 96 well plates (PrimeSurface® 3D culture, S-BIO, Japan) with a density range of 60 to 90 cells per well in 200 µl of appropriate chemically defined chondrogenic medium formulation. The cells were left to form aggregates over 48 hours at 37 °C, 5% CO2 and 95% humidity before imaging. C8 consisted of LG-DMEM (Gibco) supplemented with 1% antibiotic-antimycotic (100 units/mL penicillin, 100 mg/mL streptomycin and 0.25 mg/mL amphotericin B), 1 mM ascorbate-2 phosphate, 100 nM dexamethasone, 40 µg/mL proline, 20 µM of Rho-kinase inhibitor Y-27632 (Axon Medchem), ITS + Premix Universal Culture Supplement (Corning) (including 6.25 µg/mL insulin, 6.25 µg/mL transferrin, 6.25 µ/mL selenious acid, 1.25 µg/mL bovine serum albumin (BSA), and 5.35 µg/mL linoleic acid), 100 ng/mL BMP-2 (INDUCTOS®), 100 ng/mL GDF-5 (PeproTech), 10 ng/mL TGF-*β* 1 (PeproTech), 1 ng/mL BMP-6 (PeproTech) and 0.2 ng/mL FGF-2 (PeproTech). TGF^−^ consisted of the same supplements as C8 excluding TGF-*β* 1. ROCKi^−^ consisted of the same supplements as C8 excluding Y-27632. Bleb/ROCKi^+^ consisted of the same supplements as C8 with 100 µM instead of 20 µM Y-27632 and an additional 20 µM blebbistatin. A summary of medium compositions for the 4 studied conditions is provided in SI Table 1.

### Brightfield imaging of spheroid fusion

Two spheroids (48h from seeding) were transferred per well of the U-bottom Ultra-low Attachment 96 well plate. To minimize variability, 12 replicates per condition were imaged simultaneously in a single well plate. After transfer, the plate was immediately placed in an OKOlab Top Stage Incubator with OKO-Air-Pump-BL, OKOlab 2GF-Mixer, and OKOlab T unit controller, all operated via an OKOlab Touch set at a temperature of 37°C and a CO2 concentration of 5%. Wells containing two spheroids were selected and imaged with an Olympus IX53 microscope, SC30 Colour Camera (Olympus), UPLFLN 10× magnification objective for 24 h with 10 min interval. The cellSens Dimension (Olympus) software was used for image acquisition. This procedure performed for 5 independent repeats. Images were masked using scikit-image library in Python and Image J [45].

### Fusion shape analysis

Fusion geometry was parameterized as sin^2^(*θ*) = (*x/R*)^2^, with spheroid raidus *R* (assuming equal sizes) and contact radius *x*, both evolving over time, see Fig. 1(b). The following analysis was performed on each frame of the time-lapse microscopy sequence or active foam simulation. 1) Masks were smoothed by dilating and eroding with a kernel size of 3. For two large objects (indicating no contact), outer contour pixels were extracted from both. For a single large object (indicating fusion), outer pixels of that object were extracted. 2) A double circle was fitted by searching for the center of mass points (*x*_*c*,1_, *y*_*c*,1_) and (*x*_*c*,2_, *y*_*c*,2_) that minimize the squared error on the distance from each contour pixel to the closest center of mass. For two separate objects, the center of mass was calculated by averaging the pixel positions. For a single object, an ellipse was fitted to the outer contour and the initial guess was set at -0.5 and +0.5 along the major axis of the ellipse. The previous time step was considered as an extra initial guess and the result with the lowest converged cost function was selected. 4) Upon convergence, the spheroid radius was determined from *d*_*i*_ = min(*d*_*i*,1_, *d*_*i*,2_). Using the circle centers and radius, the intersection points were calculated and subsequently the contact radius *x*, which is half the distance between the intersection points. Frames where circle fitting failed were excluded from the time series analysis of fusion dynamics. Additionally, due to the sparsity of cells in some aggregates and potential distortion of contours caused by loosely attached or detached cell clusters, or the presence of pronounced blebbing, the entire time series of fusion was discarded from the dataset.

### Visco-elastic model for tissue spheroid fusion

Fusion dynamics can be expressed in function of visco-elastic material properties by the governing equation [13]

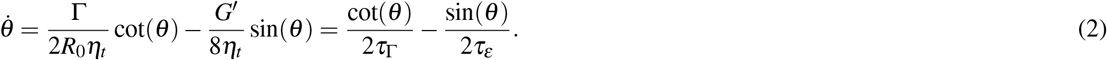

*θ* is the angle shown in Fig. 1(b), and is influenced by the tissue surface tension Γ, the radius of the spheroid before fusion *R*_0_, the apparent tissue viscosity *η*_*t*_ and the shear modulus *G*^*’*^ of the tissue. These parameters are combined into two characteristic time constants; *τ*_Γ_ is the visco-capillary time, and *τ*_*ε*_ is the visco-elastic time. The analytical solution of Eq. (2) is

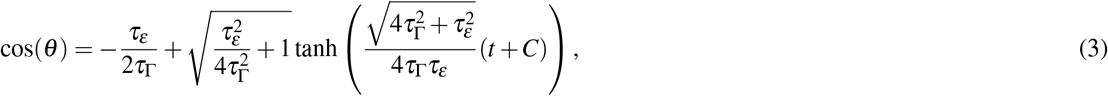

where complex number *C* can be obtained from the initial conditions *θ* (*t* = 0) = *θ*_0_

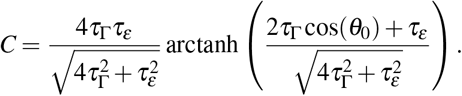

Fusion dynamics are parameterized by the time scale *τ*_*f*_ and equilibrium angle *θ*_*eq*_. Based on Eq. (3), we define *τ*_*f*_ as:

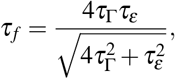

As *τ*_*ε*_→ ∞, i,e., there is no elasticity, and we obtain *τ*_*f*_ = 4*τ*_Γ_, When dominated by elasticity, i.e. 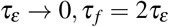. Resubstituting *τ*_*ε*_ = 4*η*_*t*_*/G*^*’*^ and *τ*_Γ_ = *η*_*t*_*a*_0_*/*Γ, we obtain

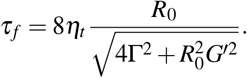

The equilibrium angle is

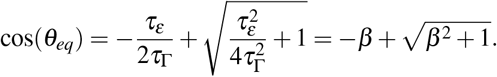

Hence, Eq. 3 can be written as

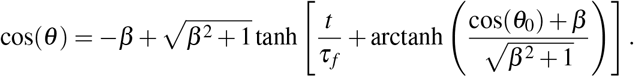

### Confocal microscopy imaging

Aggregates were fixed in 4% paraformaldehyde solution for 1 h at room temperature and washed with PBS. Aggregates were stained with the mixture of CellBrite® Fix 640 (1:1000, #30089-T, Biotinum), 1X Alexa Fluor™ 488 phalloidin (A12379, Invitrogen), 5 µg/ml Wheat Germ Agglutinin, Alexa Fluor™ 594 in PBS with 0.3 % Triton X-100 for 1 h at room temperature. Nuclei were conterstained with 1 µg/ml DAPI for 15 min at room temperature. The aggregates were washed three times with PBS and mounted into Mowiol® 4-88 (Sigma, USA). Confocal imaging was performed with LSM 780 confocal microscope (Zeiss) with a water-immersion 25× objective lens (NA 0.8, Zeiss).

### Granular shape index

The granular shape index was defined as *GSI* = *r* (*κ*−1*/R*) where *r* is the (mean) cell radius, *κ* the curvature of the tissue spheroid, and *R* the spheroid radius. In the limit of high cohesion, the spheroid approximates a perfect sphere with radius *R*, so *κ* = 1*/R* and *GSI* = 0. Conversely, at low cohesion, with *r*≪*R*, the curvature *κ*≈ 1*/r*, resulting in *GSI*→ 1. Based on earlier experiments, the radius of hPDCs follows a Gamma distribution with shape factor *k* = 37 and scale factor *ϑ* = 0.24 µm, with average radius *r* = *kϑ* = 8.88 µm [33]. From confocal microscopy images, curvature *κ* was calculated as: 1) Slices were taken in the middle of the aggregate and contrast was adjusted to enhance contour visibility. 2) Based on a custom analysis tool in Python, we approximate a section (typically corresponding to a cell) by fitting a circle through 3 manually selected points. 3) Based on the intersection points of these circles, the intersection arc was determined which forms the full contour of the aggregate slice, see Fig. 2(b). 4) 1000 equidistant points were sampled along the contour, and binned into 360 angular bins.*κ*(*φ*) is the mean curvature in each bin. Due to heavy tails in the curvature distribution from loose cells or blebs, Fig. 2(c), *κ* is defined as the median of the binned curvature. 5) A circle is fitted through the equidistant contour points, searching the center of mass point (*x*_*c*_,*y*_*c*_) for which the distance to each boundary pixel has the lowest squared error.

### Two-photon imaging for cell tracking

hPDCs were labeled at 70% confluence in a T75 flask with 3 µg/mL Hoechst 33342 (#H3570) in 10 ml of non-additive DMEM for 20 min at 37 °C. Cells were subsequently washed, trypsinized and seeded in a U-bottom Ultra-low Attachment 96 well plate as was done for brightfield imaging. The inclusion of Hoechst 33342 did not influence cell aggregation and aggregate fusion. Labeled 48h old aggregates were transferred into a bioinert Spheroid Perfusion µ-Slide (Ibidi, 80350, Germany) and subsequently imaged on an LSM 780 confocal microscope (Zeiss) with a water-immersion 25× objective lens (NA 0.8, Zeiss). A tunable Mai Tai DeepSee Titanium-Sapphire femtosecond laser (690-1040 nm, Spectra-Physics) set at 750 nm was used for 2 photon excitation and fluorescence was detected in epi between 391-598 nm. Time-lapses were recorded with a 5 minute interval for 10h, a variable Z-stack range depending on the thickness of aggregates and a 0.29 *×*0.29*×* 1.7 µm voxel size. Images were adjusted for brightness and contrast and registered with descriptor-based registration plugins in ImageJ [46].

### Automated cell segmentation and tracking

From the confocal time-lapse series, Z-stacks were extracted for every time point and subsequently rescaled to obtain isotropic voxel dimensions. Every Z-stack was further split into 4 substacks of the Z-stack along the x- and y-direction, thus preserving the height, with a 10% overlap with neighboring substacks to reduce memory costs of computation. To account for the decay of intensity with distance from the objective, the slice-wise mean of the substacks was scaled with the overall mean intensity of the substack. The substacks were smoothed using a 3D Gaussian blur (*σ* = 1.5 µm), followed by a Frangi-Vesselness filter (*α* = 0.9, *β* = 0.9, *γ* = 1.0) to amplify blob-shaped objects. The Frangi-filtered substacks were thresholded using automatic Otsu’s thresholding to obtain a binary mask of blob-like objects of the substacks. These binary masks were recombined to the original Z-stack dimensions preserving the maximal values in overlapping regions, thus re-assembling nuclei that were potentially cut due to the separation in substacks. To obtain individual objects from the binary mask of the Z-stack, the binary mask was segmented using a watershed algorithm seeded from local maxima in the intensity signal. Foreground and background objects were separated by using Otsu’s threshold on the distribution of mean intensities of all segmented objects. Finally, to select the segmented nuclei, only large objects were retained by Otsu’s filtering the volume of the foreground objects extracted from all the Z-stacks originating from a single time-lapse experiment. The tracking over time of the segmented nuclei was automatically performed using LapTrack [47]. Frame-to-frame linking of nuclei was performed by solving a Linear Assignment Problem (LAP) with the cost function being the squared Euclidean distance between the centroids of the nuclei in successive frames. The cut-off distance for linking nuclei was set to *d*_*c*_ = 15 µm. Merging and splitting of tracks was not allowed.

### Relative diffusion measurement

The squared relative displacement was calculated between between cell pairs at various lag times (*τ >* 0). From this, the time and ensemble average was

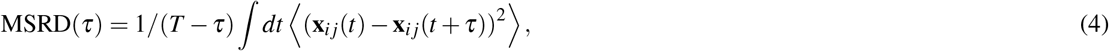

with **x**_*ij*_(*t*) the relative displacement between cells *i* and *j*, **x**_*ij*_(*t*) = **x**_*i*_(*t*) −**x** _*j*_(*t*). In simulations, pairs at time *t* are obtained based on direct contacts between cells. In experiments, a Delaunay triangulation on the center of mass of the segmented nuclei was performed and only connections shorter than 3.5*r* were retained as pairs.

### 3D active foam model

The computational cell model, based on previous approaches [25, 48, 49, 50], portrays tissues as foam-like materials. Each cell *i* is modeled as a volume-conserving, pressurized “bubble”, characterized by a thin, viscous shell with plane viscosity *η*. This represents the cell’s acto-myosin cortex-membrane complex. Cells possess polarity 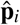,which diffuses as 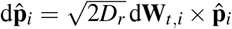,with persistence time *τ*_*p*_ = 1*/D*_*r*_ and standard Wiener process **W**_*t*_ To model active cell migration, an outward migration pressure is exerted on the cell surface as 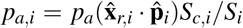,where *p*_*a*_ is the active pressure, 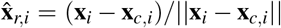is the relative position with respect to the cell center **x**_*c,i*_, *S*_*c,i*_ is the total cell-cell contact area and *S*_*i*_ is the total cell area. To preserve internal force balance, the internal reaction traction 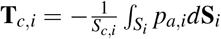 is applied on contacting surfaces. Given plane tangent vectors 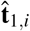 and 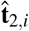, the resulting in-plane stress is ***σ*** _*i*_ = *η*∇**u**_*i*_ + *γ*_*s*_**I**, with in-plane velocity 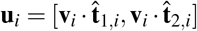..Note that for the viscous stress we have assumed irrotational Stokes flow. The net surface tension is

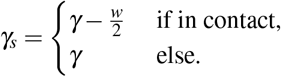

We define the cortical relaxation time *τ*_*γ*_ = *γ/η* and the relative cell-cell tension is *α* = 1−*w/γ*. The in-plane force balance for surface *i* in contact with surface *j* reads

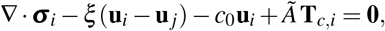

with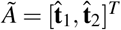, medium damping *c*_0_, and cell-cell friction *ξ*, setting the hydrodynamic length-scale *L*_*η*_ = (*η/ξ*)^1*/*2^. The out-of-plane pressure balance is

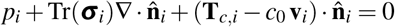

with outward normal ň. The total pressure is *p*_*i*_ = *p*_*K,i*_ + *p*_*a,i*_ + *p*_*c,i*_, with *p*_*K,i*_ the bulk pressure for volume conservation and *p*_*c,i*_ the normal contact pressure, which for overlap distance *δ*_*ij*_ is *p*_*c,i*_ = *w/h*_0_−*wδ*_*ij*_*/h*^2^, with effective range of interaction *h*_0_. To simulate multicellular collections, a particle-based computational model is adopted where the cortex-membrane complex is represented by a triangulated surface mesh, with nodal positions serving as degrees of freedom. Details of the numerical discretization and computational implementation of this 3D foam model are provided in the Supplementary Note.

### Parametric generation of virtual tissue spheroids

A library of virtual tissue spheroids was created for combinations of cell mechanical properties, but lacking cell motility. We generated *N* spherical cells with radii from a Gamma distribution, uniformly distributed within a spherical volume of radius 2*N*^1*/*3^*r*_*i*_, with *r* = 8.88 µm as the reference. The cells were enclosed by a reflective sphere with radius 2*N*^1*/*3^*r* + 2*r*. The domain’s radius linearly decreased over time *τ*_init_ = 10*τ*_*γ*_ *N*^1*/*3^ to allow relaxation. A centripetal force **F**_*cp*_(*t*) = −*F*_*cp*_ exp(−5*t/τ*_init_)**e**_*r*_ was applied to cells outside the final boundary for *t* ≤ *τ*_init_*/*5. Cells were then allowed to relax without compression for an additional *τ*_init_.

### Simulation setup

To simulate fusion, two spheroids are initially positioned with minimal overlap. First, two spheroids from the library are randomly selected and centered with random orientations. They are then translated along the **e**_*x*_ axis until their boundaries touch. The minimum translation required for geometric contact is computed; if it exceeds a critical threshold, the spheroids are moved further apart and the process is repeated. If it is below the threshold, the spheroids are moved to achieve contact and the iteration ends. Finally, the simulation is centered at the origin. All simulations were conducted in Python using the Mpacts particle-based simulation framework.

## Acknowledgments

This work is part of Prometheus, the KU Leuven R&D Division for Skeletal Tissue Engineering. J.V. acknowledges support from the Research Foundation Flanders (FWO), grant 11D9923N. Images were recorded on a Zeiss LSM 780 – SP Mai Tai HP DS (Cell and Tissue Imaging Cluster (CIC), supported by Hercules AKUL/11/37 and FWO G.0929.15 to P.v.d.B., University of Leuven. H.S. acknowledges support from MSCA4Ukraine fellow (AvH ID): 101101923. This work was supported by Interne Fondsen KU Leuven/Internal Funds KU Leuven grant numbers C24M/22/058. B.S. acknowledges support from the Research Foundation Flanders (FWO), grant 12Z6118N, and KU Leuven internal funding C14/18/055.

---

**Extended Data Fig. 1:**
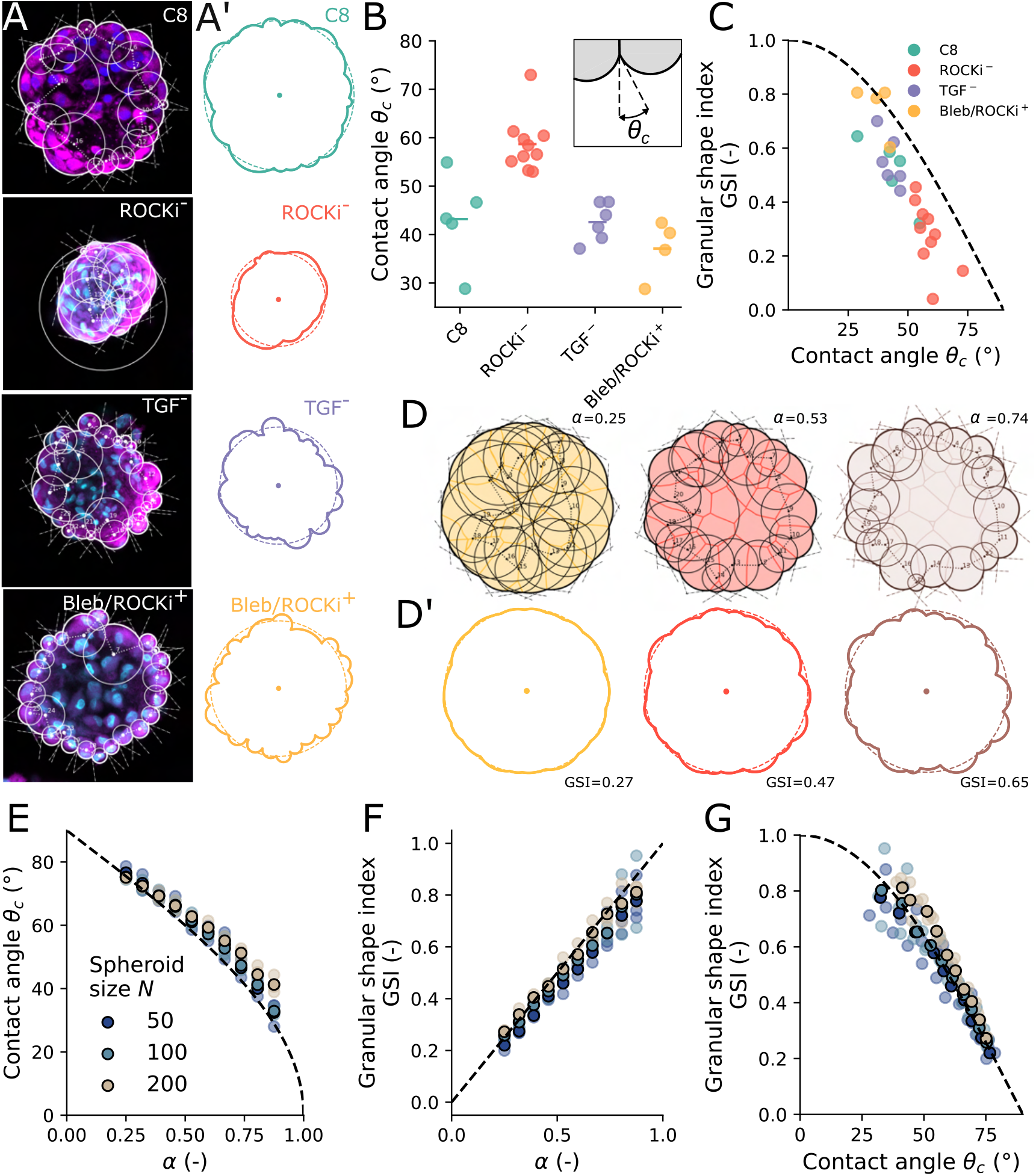
Granular shape index is a measure of relative cell-cell tension. **A**: Confocal microscopy slices of representative spheroids in conditions C8, ROCKi^−^, TGF^−^ and Bleb/ROCKi^+^, with DAPI staining (nucleus), annotated with fitted contour circles, and **A’**, extracted spheroid contour with fitted spheroid radius. **B**: Extracted contact angle *θ*_*c*_ (inset) in 4 conditions, based on the contact angle between individual cells at the aggregate contour. Each data point represents the median contact angle from a single spheroid, with the horizontal bar indicating the average across *n* spheroids in each condition. Measurements are biased towards lower contact angles due to difficulties in distinguishing contacts between neighboring cells with low curvature (compare a and d) and the presence of bleb-like protrusions in otherwise smooth aggregates. **C**: Granular shape index (GSI) in function of extracted contact, with theoretical scaling *GSI* = cos(*θ*_*c*_) as a dashed line. **D**: Slices of simulated tissue spheroids at varying *α*, annotated with fitted contour circles, and **D’**, extracted spheroid contour with fitted spheroid radius, annotated with the corresponding *GSI*. **E**: In simulated spheroids of varying size *N*, where *θ*_*c*_ can be accurately determined, contact angle measures relative cell-cell tension *α*. The dashed line shows *θ*_*c*_ = arccos(*α*). **F**: Granular shape index measures *α* in simulated tissue spheroids. The dashed line shows *GSI* = *α*. **G**: *GSI* closely follows the theoretical scaling, *GSI* = cos(*θ*_*c*_), in simulated tissue spheroids.

**Extended Data Fig. 2:**
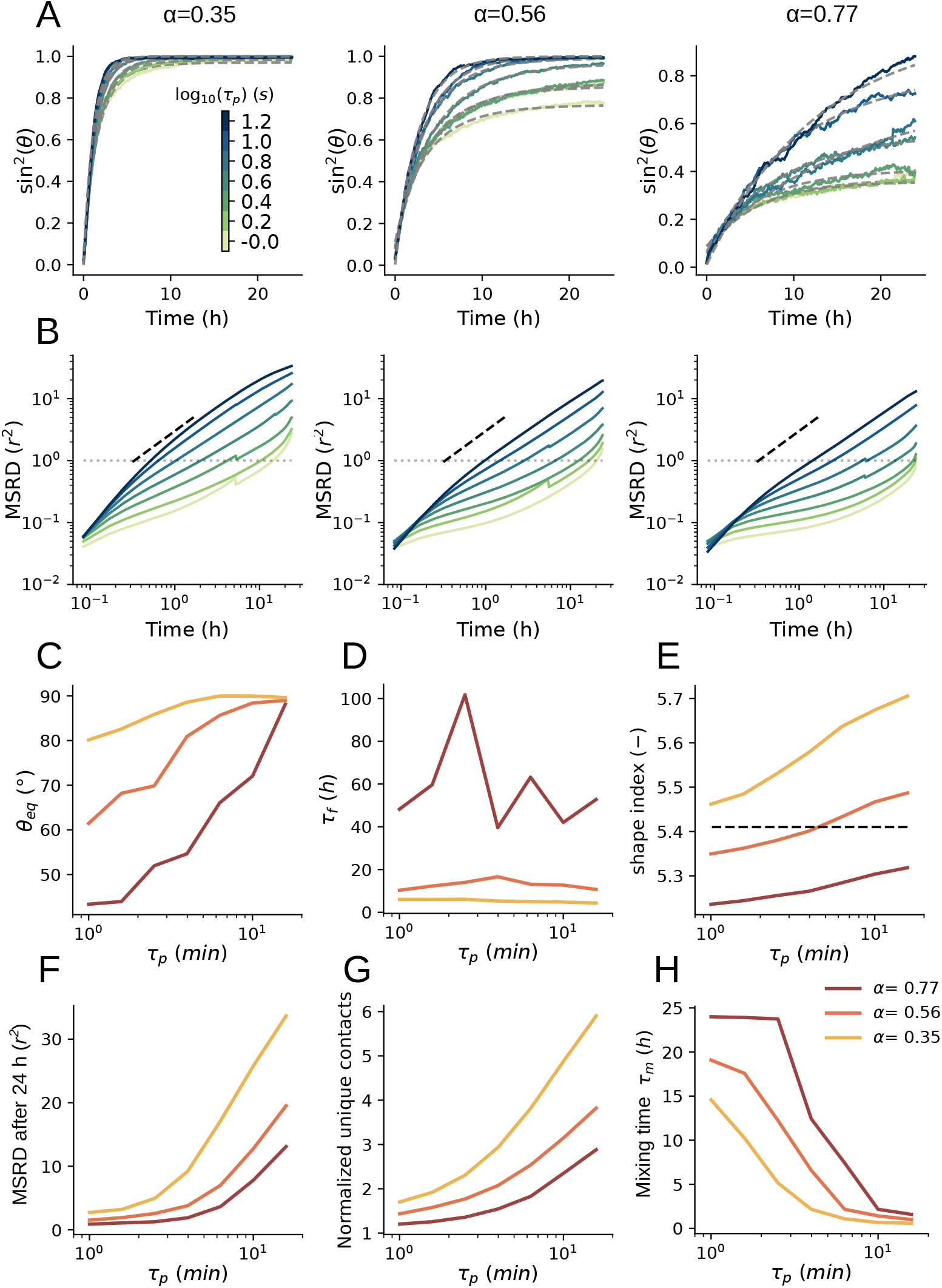
Persistence time affect fusion dynamics and tissue fluidity. **A**: Fusion angle in function of time for simulated spheroids with varying persistence time *τ*_*p*_, at three levels of relative cell-cell tension *α*. Decreased *τ*_*p*_ leads to incomplete fusion. Fitted curves of Eq. (1) are shown with fitted parameters in c,d. **B**: Mean squared relative displacement (MSRD) in function of time for varying *τ*_*p*_ and *α*, with horizontal dashed line indicating *MSRD* = *r*^2^ and black dashed line indicating diffusive scaling. Decreasing *τ*_*p*_ induces jamming. **C**: Equilibrium fusion angle *θ*_*eq*_ increases with *τ*_*p*_. **D**: *τ*_*p*_ has no pronounced effect on fusion time *τ*_*f*_. **E**: Shape index increases with *τ*_*p*_, indicating larger cell deformations and tissue fluidization. The dashed line indicates prediction of a critical shape index at 5.41 from 3D vertex models [39]. **F**: MSRD after 24 h increases with *τ*_*p*_. **G**: Normalized unique contacts, indicating frequency of cell rearrangements, increases with *τ*_*p*_. Note the similarities between f and g. **H**: Microscopic mixing time, defined as the lag-time for which *MSRD* = *r*^2^ and capped at 24h, decreases with *τ*_*p*_.

**Extended Data Fig. 3:**
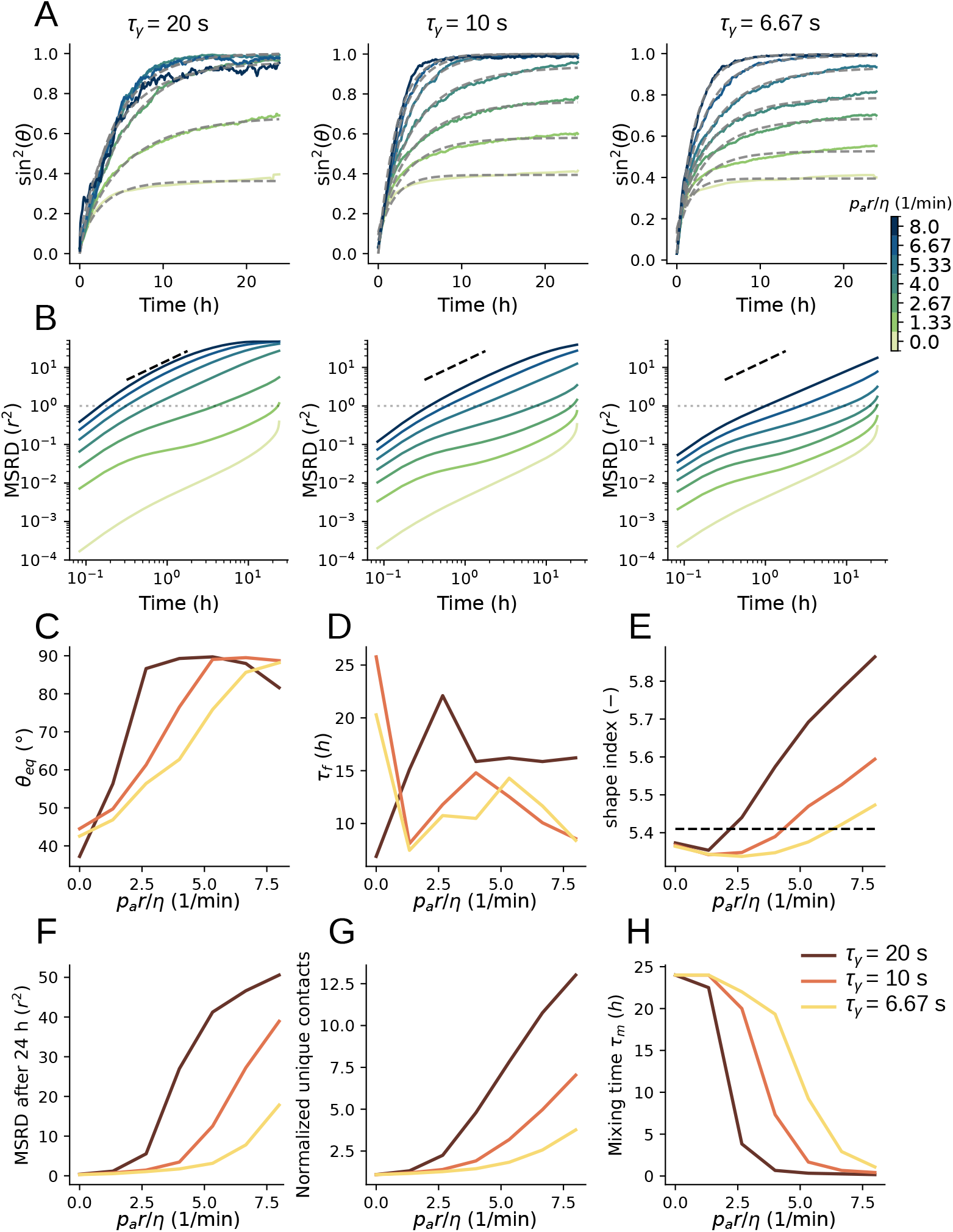
Surface tension affects fusion dynamics by promoting jamming. **A**: Fusion angle in function of time for simulated spheroids with varying rescaled motility 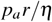 at three levels of surface tension (cortex tension at the cell-medium interface), parameterized by cortical relaxation time *τ*_*γ*_ = *η/γ*. Decreased *τ*_*γ*_ leads to incomplete fusion. Fitted curves of Eq. (1) are shown with fitted parameters in c,d. **B**: Mean squared relative displacement (MSRD) in function of time for varying *τ*_*γ*_ and 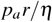, with horizontal dashed line indicating *MSRD* = *r*^2^ and black dashed line indicating diffusive scaling. Decreasing *τ*_*γ*_ reduces relative displacements. **C**: Equilibrium fusion angle *θ*_*eq*_ increases with *τ*_*γ*_. **D**: Fusion time *τ*_*f*_ decreases with *τ*_*γ*_ at low cell motility, but increases at high cell motility. **E**: Shape index increases with *τ*_*γ*_, indicating larger cell deformations and tissue fluidization. The dashed line indicates prediction of a critical shape index at 5.41 from 3D vertex models [39]. **F**: MSRD after 24 h increases with *τ*_*γ*_. **G**: Normalized unique contacts, indicating frequency of cell rearrangements, increases with *τ*_*γ*_. Note the similarities between f and g. **H**: Microscopic mixing time, defined as the lag-time for which *MSRD* = *r*^2^ and capped at 24h, decreases with *τ*_*γ*_.

**Extended Data Fig. 4:**
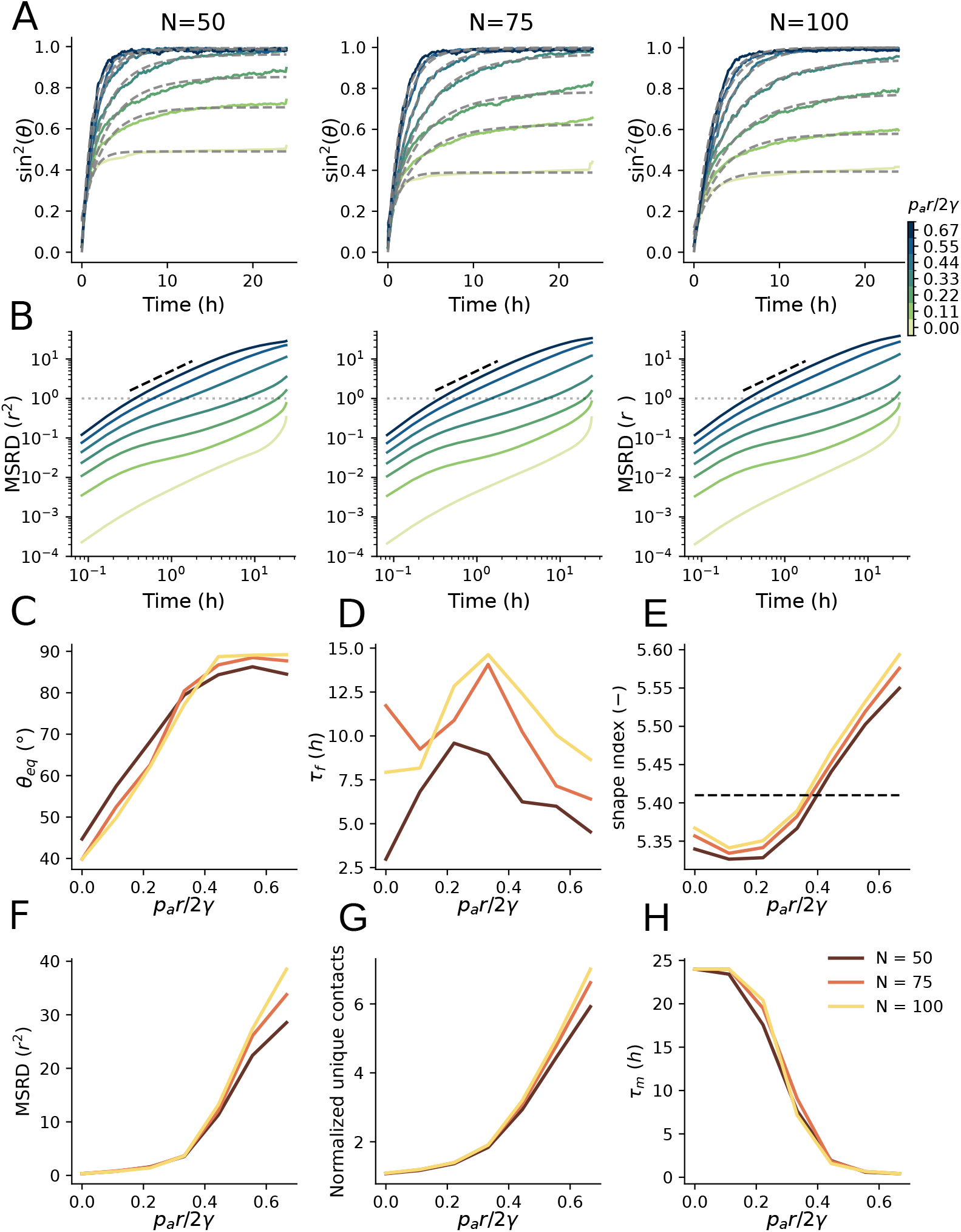
Weak size effects on fusion dynamics and tissue fluidity for small spheroids. **A**: Fusion angle in function of time for simulated spheroids with varying motility 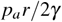, at three spheroid sizes, with number of cells *N* = 50, 75, 100. Increased size weakly favors incomplete fusion. **B**: Mean squared relative displacement (MSRD) in function of time for varying *N* and 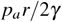,with horizontal dashed line indicating *MSRD* = *r*^2^ and black dashed line indicating diffusive scaling. There is no pronounced effect of spheroid size on relative displacements. **C**: At low motility, smaller spheroids show more complete fusion. At high motility, decreased spheroid size leads to *θ*_*eq*_ *<* 90^°^, as active surface fluctuations are more pronounced at small *N*. In the studied range of spheroid sizes, these effects are weak. **D**: Fusion time increases with spheroid size. **E**: Shape index, indicating cell deformations, weakly increases with spheroid size. The dashed line indicates prediction of a critical shape index at 5.41 from 3D vertex models [39]. **F**: No pronounced effect of spheroid size on the MSRD after 24 h. **G**: Normalized unique contacts, indicating frequency of cell rearrangements, do not strongly depend on spheroid size. **H**: Microscopic mixing time, defined as the lag-time for which *MSRD* = *r*^2^ and capped at 24h, does not depend on spheroid size.

**Extended Data Fig. 5:**
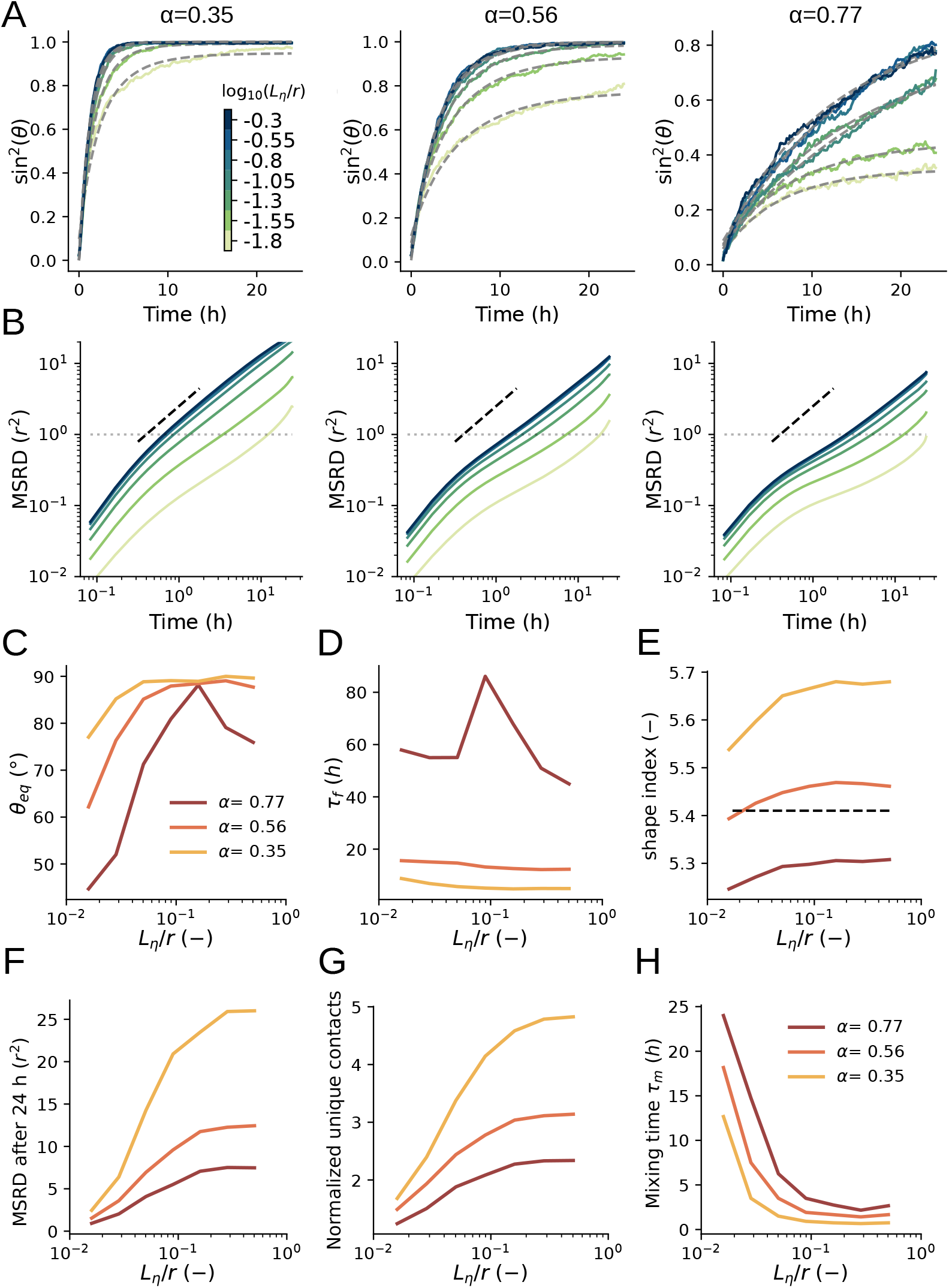
Cell-cell friction induces arrested fusion. **A**: Fusion angle in function of time for simulated spheroids with varying cell-cell friction, at three levels of relative cell-cell tension *α*. Friction is parameterized through the hydrodynamic length-scale *L*_*η*_ = (*η/ξ*)^1*/*2^, with cortex viscosity *η* and cell-cell friction constant *ξ* (High values of *L*_*η*_ correspond to low cell-cell friction). Cell-cell friction induces arrested fusion. **B**: Mean squared relative displacement (MSRD) in function of time for varying *L*_*η*_ and *α*, with horizontal dashed line indicating *MSRD* = *r*^2^ and black dashed line indicating diffusive scaling. Cell-cell friction decreases MSRD and promotes jamming. **C**: Cell-cell friction leads to incomplete fusion. At low friction and large *α*, the spheroid is an agitated granular pile with equilibrium fusion angle *θ*_*eq*_ *<* 90^°^. **D**: Cell-cell friction weakly increases fusion time *τ*_*f*_. **E**: Shape index weakly decreases with cell-cell friction. At high friction and low cell-cell tension, jamming (see f,g), is induced for elongated cells (shape index ¿ 5.41), indicating that jamming induced by cell-cell friction is a distinct mechanism from shape driven rigidity transitions. The dashed line indicates prediction of a critical shape index at 5.41 from 3D vertex models [39]. **F**: Cell-cell friction sharply decreases the MSRD after 24h. **G**: Normalized unique contacts, indicating frequency of cell rearrangements, sharply decrease with increased cell-cell friction **H**: Mixing is strongly reduced by increasing cell-cell friction, as indicated by the microscopic mixing time, defined as the lag-time for which *MSRD* = *r*^2^ (capped at 24h).

## Supplementary Figures

**Supplementary Table 1:**
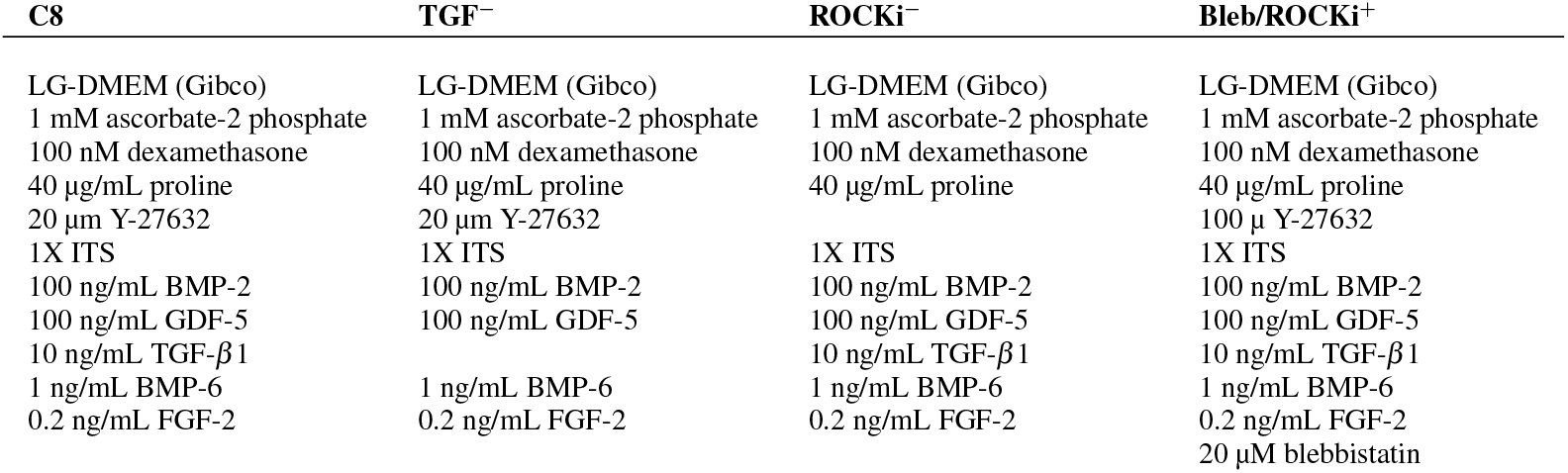
Different medium composition used in fusion experiments of hPDC tissue spheroids in conditions C8, TGF^−^, ROCKi^−^ and Bleb/ROCKi^+^

| C8 | TGF<sup>-</sup> | ROCKi<sup>-</sup> | Bleb/ROCKi<sup>+</sup> |
| --- | --- | --- | --- |
| LG-DMEM (Gibco) | LG-DMEM (Gibco) | LG-DMEM (Gibco) | LG-DMEM (Gibco) |
| 1 mM ascorbate-2 phosphate | 1 mM ascorbate-2 phosphate | 1 mM ascorbate-2 phosphate | 1 mM ascorbate-2 phosphate |
| 100 nM dexamethasone | 100 nM dexamethasone | 100 nM dexamethasone | 100 nM dexamethasone |
| 40 µg/mL proline | 40 µg/mL proline | 40 µg/mL proline | 40 µg/mL proline |
| 20 µM Y-27632 | 20 µM Y-27632 |  | 100 µ Y-27632 |
| 1X ITS | 1X ITS | 1X ITS | 1X ITS |
| 100 ng/mL BMP-2 | 100 ng/mL BMP-2 | 100 ng/mL BMP-2 | 100 ng/mL BMP-2 |
| 100 ng/mL GDF-5 | 100 ng/mL GDF-5 | 100 ng/mL GDF-5 | 100 ng/mL GDF-5 |
| 10 ng/mL TGF-β1 |  | 10 ng/mL TGF-β1 | 10 ng/mL TGF-β1 |
| 1 ng/mL BMP-6 | 1 ng/mL BMP-6 | 1 ng/mL BMP-6 | 1 ng/mL BMP-6 |
| 0.2 ng/mL FGF-2 | 0.2 ng/mL FGF-2 | 0.2 ng/mL FGF-2 | 0.2 ng/mL FGF-2 |
|  |  |  | 20 µM blebbistatin |

**Supplementary Fig. 1.**
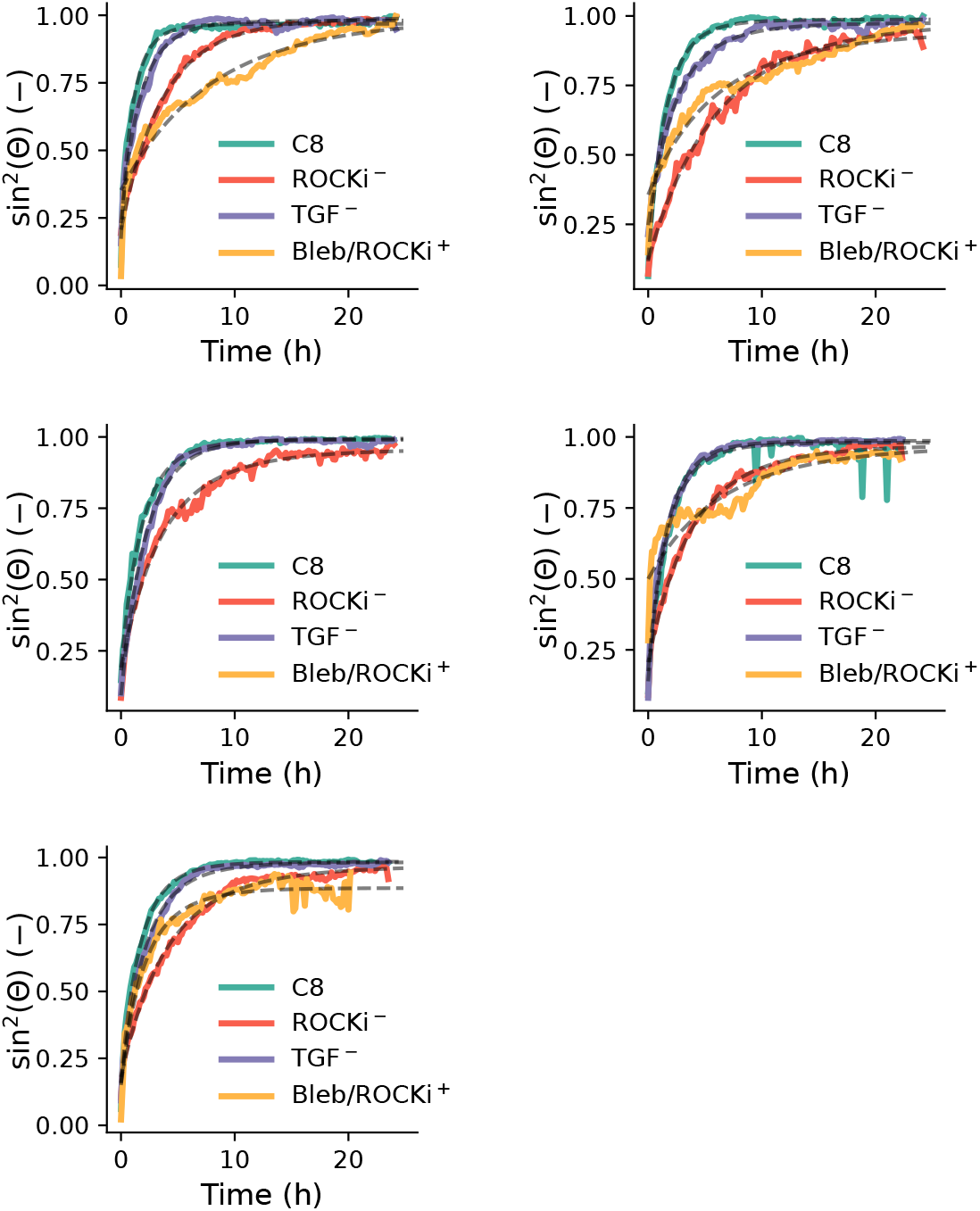
Temporal evolution of the average fusion angle for four medium compositions, together with the fit of the governing dynamics 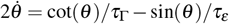. Data acquired on the same day is shown in each subplot.

**Supplementary Fig. 2.**
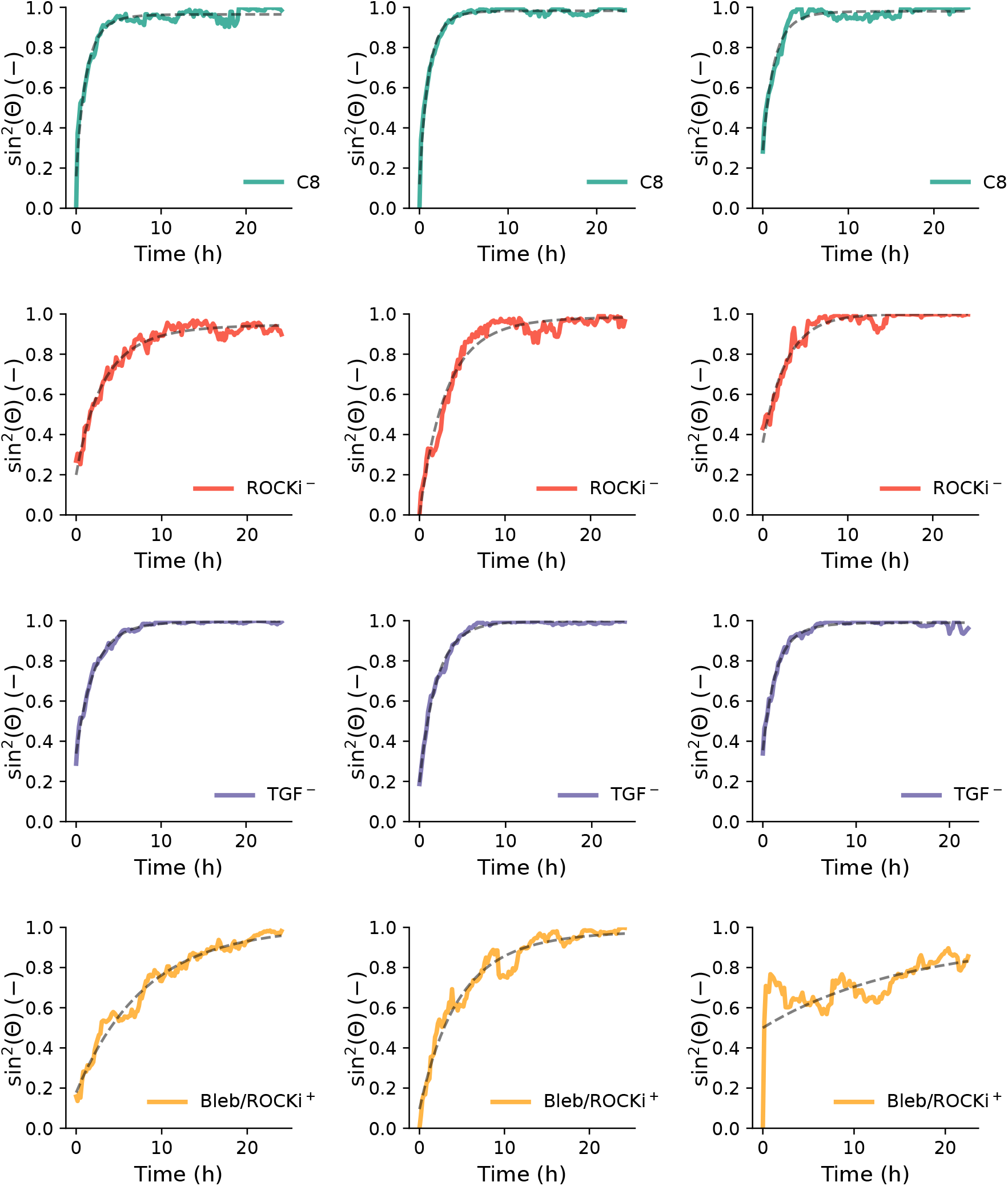
Illustration of goodness of fit on single observation of tissue spheroid fusion, as shown by the temporal evolution of the fusion angle for three individual replicas and 4 media compositions, together with the fit of the governing dynamics 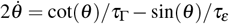.

**Supplementary Fig. 3.**
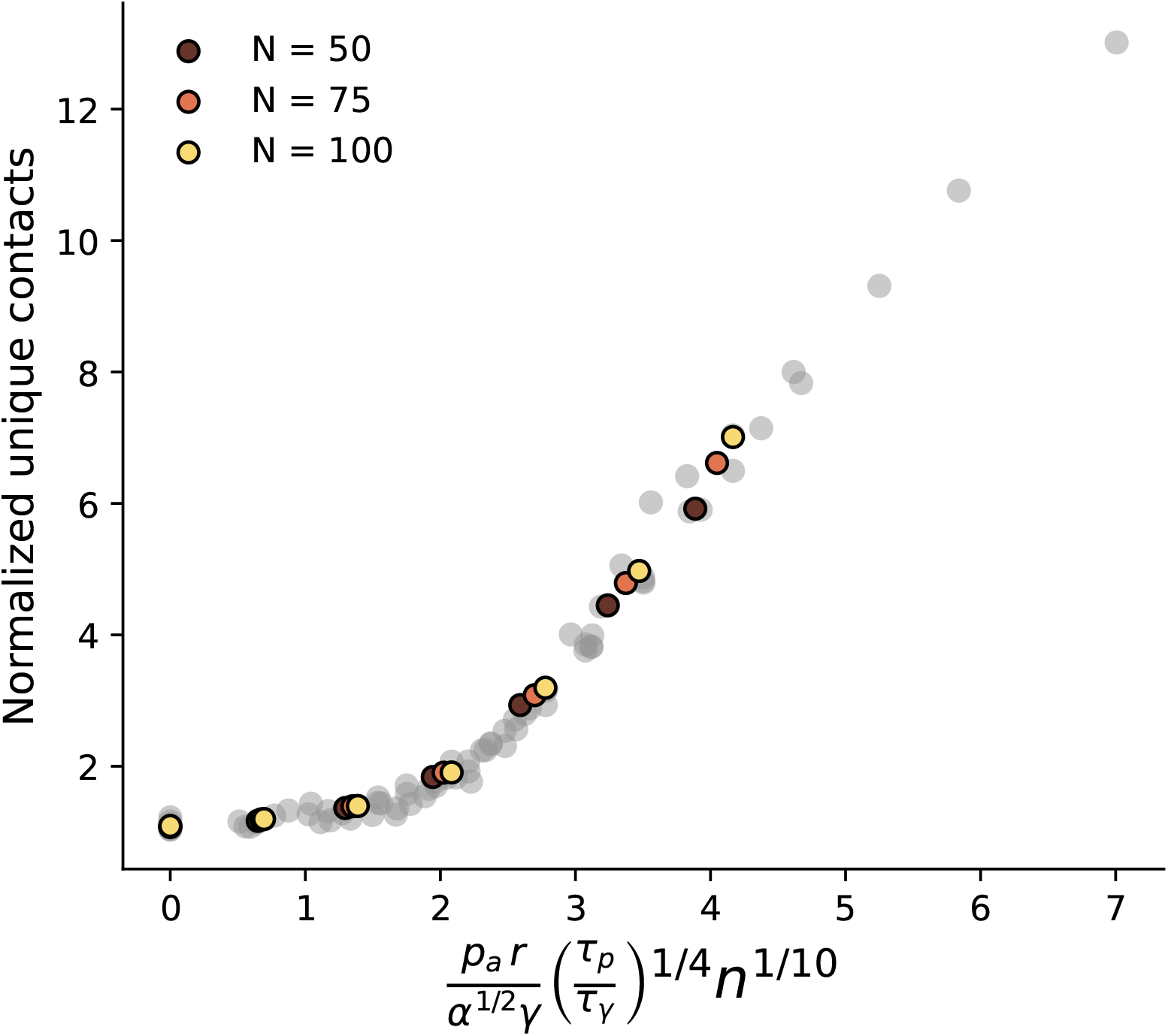
Normalized unique contacts for varying *p*_*a*_, *τ*_*p*_, and *α* (background), as well as number of cells *N* (marker color). A weak scaling (approximately ∝ *N*^1*/*10^) with cell number can be observed. (Extension of Fig. 4E)

**Supplementary Fig. 4.**
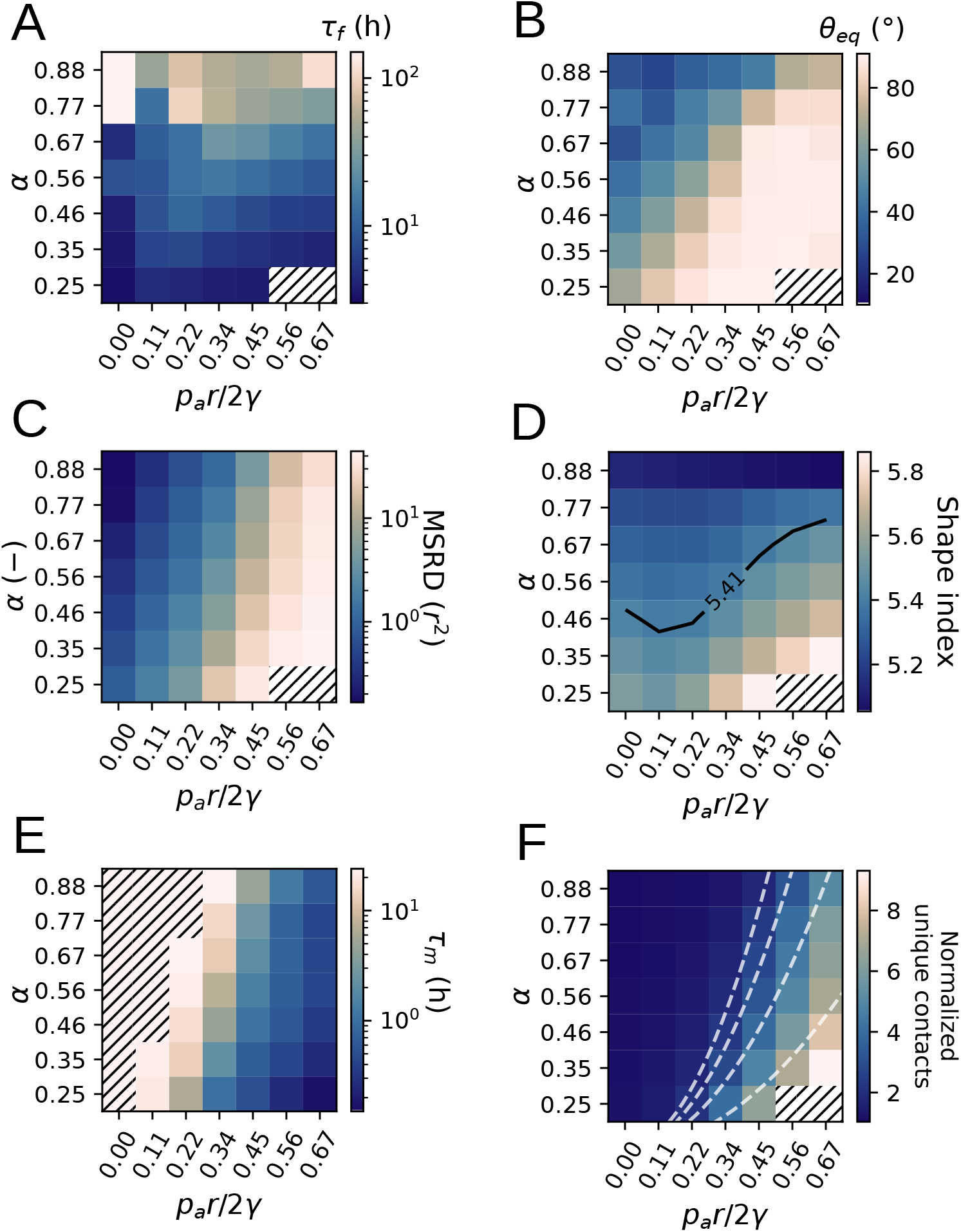
Detailed overview of the effect of active protrusion pressure *p*_*a*_ and relative cell-cell tension *α* on fusion dynamics and cell motility. **A)** Fusion time-scale *τ* _*f*_. **B)** Equilibrium fusion angle *θ*_*eq*_. **C)** Mean squared relative displacement (MSRD) after 24 hours of fusion. **D)** Shape index at the end of fusion. 3D vertex models predict a transition form jammed to fluidized at around 5.41, which is shown by the black dashed line. **E)** Microscopic mixing time which is defined as the lag-time for *MSRD* = *r*^2^. Here, the zone hatched indicates the parameter region where this threshold was not reached at the end of the 24 hour simulation. **F)** Normalized unique contacts after 24 hours of fusion. Dashed lines show the scaling 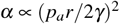. Hatched data points in (a, b, c, d, f) show simulations that failed to converge at high motility and low *α*.

## Notes

### Competing Interest Statement

The authors have declared no competing interest.

