## Supplementary Note for "Active foam dynamics of tissue spheroid fusion"

### DEFORMABLE CELL MODEL

In this supplementary note, we present the details of the deformable cell model, which is akin to the work of Cuvelier et. al. [1], where viscous cells are modeled using a particle-based representation. The tissue is treated as an active foam where each cell behaves as a pressurized "bubble" with surface tension, adhesion and internal pressure. The acto-myosin cortex of a cell is modeled as a viscous, tensed shell, approximated by a triangulated surface mesh with effective thickness $t$, and nodal positions $\boldsymbol{x}_i$ as the primary (translational) degrees of freedom. Forces acting on the nodes are categorized into three distinct types based on their origin. The subsequent sections will discuss the different force contributions and their implementation.

#### Surface tension

Tension generated in the cortex results from acto-myosin contractility, which is represented as the effective free-surface tension $\gamma$ in the shell model. Moreover, the in-plane contribution of interfacial adhesion energy $w$ generates a negative surface tension in contacting surfaces. Hence, the net surface tension $\gamma_s$ on a given surface element is

$$\gamma_s = \begin{cases} \gamma - \frac{w}{2}, & \text{for contacting surfaces and,} \\ \gamma & \text{for free surfaces.} \end{cases} \tag{1}$$

The factor $1/2$ for the adhesion energy is included because in our model $w$ represents the 'net work of adhesion' between two contacting surfaces. It should be noted that, in our model, there is no fundamental difference between adhesion and (reduction in) interfacial tension. Hence, while the underlying physical mechanism that governs contact angles in biological systems is a reduction in contractility [2], the net physical effect is an adhesive tension. Both the effects of adhesion and reduced interfacial tension are captured in the relative cell-cell tension,

$$\alpha = 1 - \frac{w}{\gamma}, \tag{2}$$

which uniquely parameterizes the contact angle $\theta_c$ between two adjacent cells, $\cos(\theta_c) = \alpha$. Using the derivation of Fedosov et al. [3], the direct force contribution of surface tension $\gamma_s$ of node $i$ is given by

$$\boldsymbol{F}_i^{\text{act}} = \frac{\gamma_s}{2}(\boldsymbol{x}_k - \boldsymbol{x}_j) \times \hat{\boldsymbol{n}}_A, \tag{3}$$

where $i, j, k$ are nodes of a single triangle $A = (ijk)$.

#### Internal pressure

Since the cytoplasm is assumed to be incompressible, active volume control is implemented via a cytoplasmic pressure. We assume the equilibrium volume $V_l^*$ of a cell $l$ is constant and implemented a proportional-integral (PI) pressure controller to enforce this. The cytoplasmic pressure $P_l^{\text{cyt}}(t)$ due to the volume controller is then

$$P_l^{\text{cyt}}(t) = -K\varepsilon_l(t) - \frac{K}{\tau_K}\int_0^t \mathrm{d}s\, \varepsilon_l(s), \tag{4}$$

for a cell with volumetric strain $\varepsilon_l(t) = (V_l(t) - V_l^*)/V_l^*$. The total pressure acting on a cell

$$P_l(t) = \frac{2\gamma}{r_l} + P_l^{\text{cyt}}(t), \tag{5}$$

includes an additional offset pressure $2\gamma/r_l$ to balance the acto-myosin contractility in the cortex. This ensures cells are in mechanical equilibrium at the start of the simulations. The force acting on triangle $A$ due to internal pressure $P_l(t)$, is obtained by multiplying with the triangle area vector $\boldsymbol{S_A} = S_A\hat{\boldsymbol{n}}_A$. Given the pressure is constant over the triangle surface, the force per triangle is

$$\boldsymbol{F}_A^{\text{cyt}}(t) = \boldsymbol{S}_A\, P_l(t). \tag{6}$$

These volume forces are then equally distributed to the nodes $(ijk)$.

### Cell migration

Cell migration is modeled as a protrusive pressure at the leading cell edge and a contractile pressure at the trailing edge, resembling mesenchymal migration by actomyosin polymerisation and cell contraction, respectively. The pressure on node $i$ in cell $l$ is modeled as

$$P_i^a = p_a \frac{S_{c,l}}{S_l} \frac{\boldsymbol{x}_i - \boldsymbol{x}_{c,l}}{||\boldsymbol{x}_i - \boldsymbol{x}_{c,l}||} \cdot \hat{\boldsymbol{p}}_l \tag{7}$$

where $p_a$ is the active pressure, $S_{c,l}$ is the cell-cell contact area of cell $l$ and $S_l$ is the total cell area, $x_{c,l}$ is cell center and $\hat{\boldsymbol{p}}_l$ is the cell polarity. The active force on node $i$ is

$$\boldsymbol{F}_i^{\text{act}} = p_i^a \hat{\boldsymbol{n}}_i. \tag{8}$$

We assume that the polarity $\hat{\boldsymbol{p}}_l$ undergoes rotational diffusion

$$\mathrm{d}\hat{\boldsymbol{p}}_l = \sqrt{2D_r}\,\mathrm{d}\boldsymbol{W}_{t,l} \times \hat{\boldsymbol{p}}_l \tag{9}$$

with persistence time $\tau_p = 1/D_r$ and standard Wiener process $\boldsymbol{W}_{t,l}$ [4]. The internal force balance is preserved by distributing a reaction traction force over the cell contact area

$$\boldsymbol{F}_i^{\text{react}} = \begin{cases} -\frac{S_i}{S_{c,l}} \sum_{j \in l} P_j^a S_j \hat{\boldsymbol{n}}_j & \text{for contacting surfaces and,} \\ \boldsymbol{0} & \text{else.} \end{cases} \tag{10}$$

This guarantees that $\sum_i \boldsymbol{F}_i^{\text{act}} + \boldsymbol{F}_i^{\text{react}} = \boldsymbol{0}$, ensuring conservation of internal momentum.

### Medium damping

Medium drag is included as

$$\boldsymbol{F}_i^{\text{drag}} = -\boldsymbol{\Gamma}_{ii}^m \boldsymbol{v}_i,$$

where

$$\boldsymbol{\Gamma}_{ii}^m = \frac{3\eta_m}{2r} S_i \boldsymbol{I}, \tag{11}$$

with apparent medium viscosity $\eta_m$. While $\eta_m$ could be interpreted as the viscosity of the cell surrounding (i.e., water), this contribution is negligible compared to the long-time viscous behavior of multicellular systems. Hence, $\eta_m$ should be interpreted as an artificial medium damping that is included to improve the numerical stability of the integration scheme [5]. In this work, $\eta_m$ is chosen to be much lower than other relevant viscosities in the system to ensure that the precise value of $\eta_m$ does not affect the outcome of the simulations.

### Cortex viscosity

#### *Planar extensional viscosity*

A significant contribution to energy dissipation arises from 2D cortex viscosity (units of N·s/m), which we assume arises at long timescales due to remodeling of the acto-myosin cortex [6]. The viscous damping force between two connected nodes of the triangular mesh is given by

$$\boldsymbol{F}_{ij}^{\text{visc}} = \boldsymbol{\Gamma}_{ij}^c((\hat{\boldsymbol{n}}_{ij} \cdot (\boldsymbol{v}_j - \boldsymbol{v}_i))\hat{\boldsymbol{n}}_{ij} + (\hat{\boldsymbol{t}}_{ij} \cdot (\boldsymbol{v}_j - \boldsymbol{v}_i))\hat{\boldsymbol{t}}_{ij}) + \varepsilon_n(\hat{\boldsymbol{b}}_{ij} \cdot (\boldsymbol{v}_j - \boldsymbol{v}_i))\hat{\boldsymbol{b}}_{ij})), \tag{12}$$

with friction element

$$\boldsymbol{\Gamma}_{ij}^c = \frac{\eta}{\sqrt{3}},$$

and unit vectors

$$\hat{\boldsymbol{n}}_{ij} = \frac{\boldsymbol{x}_j - \boldsymbol{x}_i}{||\boldsymbol{x}_j - \boldsymbol{x}_i||},$$

$$\hat{\boldsymbol{b}}_{ij} = \frac{\hat{\boldsymbol{n}}_A + \hat{\boldsymbol{n}}_B}{\|\hat{\boldsymbol{n}}_A + \hat{\boldsymbol{n}}_B\|},$$

$$\hat{\boldsymbol{t}}_{ij} = \hat{\boldsymbol{b}}_{ij} \times \hat{\boldsymbol{n}}_{ij},$$

with $\hat{\boldsymbol{n}}_A$, and $\hat{\boldsymbol{n}}_B$, the normal unit vectors of the triangles making up the segment between nodes $i$ and $j$. We further set $\varepsilon_n = 1/10$ as a "numerically small" out-of-plane shear viscosity to prevent spurious out-of-plane numerical oscillations. This introduces artificial bending viscosity in the model, which is sufficiently low to not affect the results of our simulations [1].

### Wet contact friction

A viscous contact force is included to account for drag between contacting surfaces with friction constant $\xi$ (units of Pa·s/m), expressing the scaling between dissipative traction and the sliding velocity of two contacting surfaces $A$ and $B$

$$\boldsymbol{T}_{AB}^{\text{fric}} = -\xi \Delta \boldsymbol{v}_{AB}. \tag{13}$$

In the particle-based model, the contact drag force acting on node $i$ of triangle $A$ of the contact pair $(AB)$ is computed as

$$\boldsymbol{F}_{AB,i}^{\text{fric}} = \boldsymbol{\Gamma}_{AB}^{\text{fric}} \cdot \sum_{k \in B} w_{ik}^{AB} \left(\boldsymbol{v}_k - \boldsymbol{v}_i\right) \tag{14}$$

determined by a friction tensor $\boldsymbol{\Gamma}_{AB}^{\text{fric}}$ and weights $w_{ik}^{AB}$ per node $k$ of the $B$ triangle. $w_{ik}^{AB}$ are assumed to scale with the relative contribution of the nodal contact forces to the overall contact force thus

$$w_{ik}^{AB} = \frac{\left(\boldsymbol{F}_{AB,i}^{\text{adh}} + \boldsymbol{F}_{AB,k}^{\text{adh}}\right) \cdot \hat{\boldsymbol{n}}_{AB}}{6 \sum\limits_{\forall k \in B} \boldsymbol{F}_{AB,k}^{\text{adh}} \cdot \hat{\boldsymbol{n}}_{AB}}$$

is used as an approximation. $\boldsymbol{\Gamma}_{AB}^{\text{fric}}$ for a given contact area $S_{AB}$ is estimated as

$$\boldsymbol{\Gamma}_{AB}^{\text{fric}} = S_{AB} \left[\xi^{\perp} \hat{\boldsymbol{n}}_{AB} \otimes \hat{\boldsymbol{n}}_{AB} + \xi^{\parallel} \left(\boldsymbol{I} - \hat{\boldsymbol{n}}_{AB} \otimes \hat{\boldsymbol{n}}_{AB}\right)\right],$$

with normal and tangential friction coefficients $\xi^{\perp}$ and $\xi^{\parallel}$ respectively. Note that Eq. (14) can be equivalently formulated as

$$\boldsymbol{F}_{AB,i}^{\text{fric}} = \sum_{k \in B} \boldsymbol{\Gamma}_{AB,ik}^{\text{fric}} \cdot \left(\boldsymbol{v}_k - \boldsymbol{v}_i\right),$$

if we denote

$$\boldsymbol{\Gamma}_{AB,ik}^{\text{fric}} = w_{ik}^{AB} \boldsymbol{\Gamma}_{AB}^{\text{fric}}. \tag{15}$$

### Contact forces

In the mechanical cell model, we assume a fixed and uniform adhesive tension $w$ across the cell surface. The adhesive tension incorporates cell-cell adhesive interactions as well as the effect of cortical tension reduction at the cell-cell interface [2, 7, 8]. The total adhesive tension $w$ acting on interface $(AB)$ is given by $w = w_A + w_B$. A linear force, scaling with contact overlap $\delta$, was introduced to represent repulsive interactions between surfaces and ensure stable contacts. The contact potential for contact-pair $(AB)$ with contact area $S_{AB}$ is then given by

$$E_{AB}^{\text{adh}} = \left(\frac{\delta_{AB}}{h_0} - \frac{\delta_{AB}^2}{2h_0^2}\right) w S_{AB}$$

where a distinction can be made between the attractive and repulsive energy contributions respectively. The effective range of adhesion $h_0$ was incorporated by virtually translating contact surfaces along their normal direction. The contact overlap distance $\delta_{AB}$ is calculated with respect to these translated surfaces.

The resulting contact pressure $P_{AB}^{\text{adh}}$ is estimated with a linear contact pressure model where no energy is stored in deformation at equilibrium

$$P_{AB}^{\text{adh}}(\boldsymbol{x}) = k_{AB}\delta_{AB}(\boldsymbol{x}) - P_{AB}^{0},$$

contact stiffness $k_{AB}$ and adhesive pressure $P_{AB}^{0}$ need to be set. To ensure that at equilibrium ($P_{AB}^{\text{adh}} \equiv 0$) the work of adhesion $w$ is recovered, contact stiffness has to be defined as $k_{AB} = P_{AB}^{0}/h_0$ with $P_{AB}^{0} = w/h_0$. In contrast to previous work [1], it was opted to keep $k_{AB}$ constant because at low adhesion the contact between cells was insufficiently stiff leading to cells collapsing in each other. We set $k_{AB}$ based on a middle adhesion level at reference surface tension $\gamma_{\text{ref}}$ (see Table 1), $w_A = w_B = 0.4\gamma_{ref}$, and a reference effective range $h_{0,\text{ref}}$ =75 nm, hence $k_{AB} \approx 711$ kPa/nm. This implies that the effective range changes when varying adhesion i.e. $h_0 = \sqrt{w/k_{AB}}$. Nevertheless, the specific value $h_0$ does not influence simulation outcome due to the scale separation, $h_0 \ll R^{\text{cell}}$. The contact overlap distance at point $\boldsymbol{x}$ is obtained as

$$\delta_{AB}(\boldsymbol{x}) = \max\left\{0;\ 2(\boldsymbol{x} - \boldsymbol{s}_{AB}) \cdot \left(\hat{\boldsymbol{n}}_{AB} \times \hat{\boldsymbol{l}}_{AB}\right) \tan\theta\right\},$$

where $\theta$ is the contact angle, $\hat{\boldsymbol{n}}_{AB} \propto (\hat{\boldsymbol{n}}_B - \hat{\boldsymbol{n}}_A) \times \hat{\boldsymbol{l}}_{AB}$ the contact normal and $\boldsymbol{s}_{AB}$ is an arbitrary point on the intersection line defined by $\hat{\boldsymbol{l}}_{AB} \propto \hat{\boldsymbol{n}}_A \times \hat{\boldsymbol{n}}_B$, as reported by Smeets et al. [9]. The resulting contact forces and moment are obtained by integrating the total contact pressure over the oriented contact area $\boldsymbol{S}_{AB}$,

$$\boldsymbol{F}_{AB}^{\text{adh}} = \int \mathrm{d}\boldsymbol{S}_{AB}(\boldsymbol{x})\, P_{AB}^{\text{adh}}(\boldsymbol{x}). \tag{16}$$

To distribute the contact forces acting on the contact plains to the nodal contact forces $\boldsymbol{F}_{AB,i}^{\text{adh}}$, it is assumed that the nodal forces are collinear with the contact normal $\hat{\boldsymbol{n}}_{AB}$. This results in a system of linear equations per contact pair $(AB)$

$$\begin{aligned} \sum_{i\in A} \boldsymbol{F}_{AB,i}^{\text{adh}} &= -\int_{\boldsymbol{x}\in A\cap B} \mathrm{d}\boldsymbol{S}_{AB}(\boldsymbol{x})\, P_{AB}^{\text{adh}}(\boldsymbol{x}) \\ \sum_{i\in A} \left[(\boldsymbol{I} - \hat{\boldsymbol{n}}_{AB} \otimes \hat{\boldsymbol{n}}_{AB}) \cdot (\boldsymbol{x}_i - \boldsymbol{x}_{AB})\right] \times \boldsymbol{F}_{AB,i}^{\text{adh}} & \\ &= \int_{\boldsymbol{x}\in A\cap B} \mathrm{d}\boldsymbol{S}_{AB}(\boldsymbol{x}) \times (\boldsymbol{x} - \boldsymbol{x}_{AB})\, P_{AB}^{\text{adh}}(\boldsymbol{x}) \end{aligned} \tag{17}$$

which guarantees a unique solution for every $\boldsymbol{F}_{AB,i}^{\text{adh}}$. The contact point $\boldsymbol{x}_{AB}$ is defined as the geometric center of the intersection polygon $A \cap B$. [9] The integrals on the right hand side are calculated by using a numerical quadrature rule presented in [10] with 7 quadrature points covering each triangle of the triangulated intersection polygon $A \cap B$ obtained as an intersection of projected triangles on a common plane. The precise choice of the number of quadrature points has no impact on the results in the simulation, as long as adequate quadrature coordinates and weighting values are chosen. The solution for this system is presented and discussed in the work of Smeets et al. [9].

#### Equation of motion

In the overdamped cellular environment, inertial forces may be neglected. Based on the different contributions described above, the force balance for node $i$ gives

$$\boldsymbol{F}_i^{\text{act}} + \boldsymbol{F}_i^{\text{react}} + \boldsymbol{F}_i^{\text{cyt}} + \sum_{(AB):i\in A} \boldsymbol{F}_{AB,i}^{\text{adh}} = \Gamma_{mii} \cdot \boldsymbol{v}_i + \sum_j \boldsymbol{\Gamma}_{ij}^{\text{c}} \cdot (\boldsymbol{v}_i - \boldsymbol{v}_j) + \sum_{(AB):i\in A} \sum_{k\in B} \boldsymbol{\Gamma}_{AB,ik}^{\text{fric}} \cdot (\boldsymbol{v}_i - \boldsymbol{v}_k)$$

which can be abbreviated as

$$\boldsymbol{F}_i = \sum_j \boldsymbol{\Gamma}_{ij} \cdot \boldsymbol{v}_j \tag{18}$$

for a system consisting of $N$ nodes, where

$$\boldsymbol{\Gamma}_{ij} = \begin{cases} \boldsymbol{\Gamma}_{ii}^{m} + \sum_{k:k\neq i} \boldsymbol{\Gamma}_{ik}^{\text{c}} + \sum_{(AB):i\in A} \sum_{k\in B} \boldsymbol{\Gamma}_{AB,ik}^{\text{fric}}, & i = j, \\ -\boldsymbol{\Gamma}_{ij}^{\text{c}} - \sum_{(AB):i\in A,\, j\in B} \boldsymbol{\Gamma}_{AB,ij}^{\text{fric}}, & i \neq j. \end{cases}$$

The Cartesian components of the overall force vectors can be represented as a single $(3N \times 1)$ column matrix, $\mathbf{F} = [\boldsymbol{F}_1, \ldots, \boldsymbol{F}_i, \ldots, \boldsymbol{F}_n]^T$, while the friction matrices can be assembled to a single $(3N \times 3N)$ sparse symmetric positive definite friction matrix $\boldsymbol{\Gamma}$ [11], hence obtaining the linear system

$$\boldsymbol{\Gamma} \mathbf{v} = \mathbf{F}. \tag{19}$$

The conjugate gradient method (CGM) is used to efficiently solve this system for the node velocities $\mathbf{v}(t + \Delta t)$ at each iteration in a semi-implicit scheme. Positions of the nodes are subsequently updated using backward Euler integration,

$$\mathbf{x}(t + \Delta t) = \mathbf{x}(t) + \Delta t \, \mathbf{v}(t + \Delta t), \tag{20}$$

where $\mathbf{v}(t + \Delta t)$ are the projected new velocities obtained by solving Eq. 19 at time $t$ via the CGM.

#### Remeshing

Due to the viscous material properties of the cortex, mesh nodes tend to flow as no local area or length conservation is imposed. To maintain a mesh of sufficiently high quality, i.e. sufficiently high resolution near fine details of the cell at an acceptable number of degrees of freedom, we used a remeshing algorithm based on the surface operations defined by Brakke [12] using as input a threshold minimum and maximum triangle area, similar to [13]. The remeshing step is performed every 30 time steps to ensure a sufficiently high mesh quality. By remeshing, we implicitly assume that the nodes themselves do not carry 'mass' or other local information, but are merely considered as discretization points that represent the location of the surface. Similarly, there can be no conservative (energy conserving) potentials in the membrane, since no history of previous deformation is retained in the computational model [14]. However, these conditions are explicitly assumed by the viscous approximation of the cortex model.

### MODEL PARAMETERS

Table 1 provides a list of all parameter used to simulate tissue spheroid fusion with the active foam model. The table is divided into three categories: 1) Model parameters which are constant over all performed simulations; 2) Model parameters which can vary in between simulations of a parameter study. Unless mentioned otherwise, these are the parameters used in the reported results; 3) Simulation parameter, which are associated to the numerical solutions of the simulation. Only dimensions of length and time were accessible in experiments. Furthermore, no absolute sizes, but fusion angles were reported. Hence all simulation results are reported in reduced model parameters, which are adimensional or only contain units of time. These reduced parameters are provided in Table 2.

**Table 1. Reference table of model parameters**

| Parameter | Symbol | Value | Units |
|---|---|---|---|
| **Constants Deformable Cell Model** | | | |
| Mean cell radius | $r$ | 8.8 | µm |
| Cell radius Gamma distribution k-parameter | | 37 | – |
| Cell radius Gamma distribution theta-parameter | | 0.24 | µm |
| Contact stiffness | $k_{AB}$ | 711.11... | $\mathrm{kPa\,nm^{-1}}$ |
| Cortex surface tension reference | $\gamma_{\mathrm{ref}}$ | 0.5 | $\mathrm{nN\,\mu m^{-1}}$ |
| Effective fluid viscosity | $\eta_m$ | 0.4 | kPa s |
| Effective cortex viscosity | $\eta$ | 0.005 | $\mathrm{N\,s\,m^{-1}}$ |
| Characteristic relaxation time of the cell shell | $\tau_{\mathrm{shell}}$ | $\eta/(2\sqrt{3}\gamma_{\mathrm{ref}})$ | s |
| Cytosolic bulk modulus | $K$ | 0.5 | kPa |
| Cytosolic bulk modulus integration timescale | $\tau_K$ | $2\tau_{shell}$ | $s$ |
| **References for parameter studies** | | | |
| Number of cells | $N$ | $2\times100$ | – |
| Relative cell-cell tension | $\alpha$ | 0.4375 | – |
| Protrusion pressure | $p_a$ | 50 | Pa |
| Effective contact friction | $\xi$ | 0.25 | $\mathrm{kPa\,s\,\mu m^{-1}}$ |
| Cortex surface tension | $\gamma$ | 0.5 | $\mathrm{nN\,\mu m^{-1}}$ |
| Cell motility persistence time | $\tau_p$ | 10 | min |
| **Simulation parameters** | | | |
| Time step | $\Delta t$ | 0.1 | s |
| Remesh interval | $\tau_r$ | $5\tau_{shell}$ | s |
| Reference triangle area | $A_\Delta$ | $4\pi r^2/320$ | $\mathrm{\mu m^2}$ |
| Minimal triangle area | $A_m$ | $A_\Delta/3$ | $\mathrm{\mu m^2}$ |
| Maximum triangle area | $A_M$ | $3A_\Delta$ | $\mathrm{\mu m^2}$ |
| Eigen conjugate gradient solver tolerance | $\varepsilon_{CG}$ | 0.001 | – |

**Table 2. Table of reduced model parameters.** Base values of reduced model parameters (adimensional or in units of time). These values are used for all reported results, unless explicitly stated otherwise.

| Parameter | Symbol | Value | Units |
|---|---|---|---|
| Cortical relaxation time | $\tau_\gamma = \eta/\gamma$ | 10 | s |
| Persistence time | $\tau_p$ | 10 | min |
| Relative active pressure | $p_a r/2\gamma$ | 0.33 | – |
| Relative cell-cell tension | $\alpha = 1 - w/\gamma$ | 0.4375 | – |
| Hydrodynamic length | $L_\eta/r = (\eta/\xi)^{1/2}/r$ | 0.50 | – |
| Number of cells / spheroid | $N$ | 100 | – |
